# nSMase2-mediated exosome secretion shapes the tumor microenvironment to immunologically support pancreatic cancer

**DOI:** 10.1101/2024.09.23.614610

**Authors:** Audrey M. Hendley, Sudipta Ashe, Atsushi Urano, Martin Ng, Tuan Anh Phu, Xianlu L. Peng, Changfei Luan, Anna-Marie Finger, Gun Ho Jang, Natanya R. Kerper, David I. Berrios, David Jin, Jonghyun Lee, Irene R. Riahi, Oghenekevwe M. Gbenedio, Christina Chung, Jeroen P. Roose, Jen Jen Yeh, Steven Gallinger, Andrew V. Biankin, Grainne M. O’Kane, Vasilis Ntranos, David K. Chang, David W. Dawson, Grace E. Kim, Valerie M. Weaver, Robert L. Raffai, Matthias Hebrok

## Abstract

The pleiotropic roles of nSMase2-generated ceramide include regulation of intracellular ceramide signaling and exosome biogenesis. We investigated the effects of eliminating nSMase2 on early and advanced PDA, including its influence on the microenvironment. Employing the KPC mouse model of pancreatic cancer, we demonstrate that pancreatic epithelial nSMase2 ablation reduces neoplasia and promotes a PDA subtype switch from aggressive basal-like to classical. nSMase2 elimination prolongs survival of KPC mice, hinders vasculature development, and fosters a robust immune response. nSMase2 loss leads to recruitment of cytotoxic T cells, N1-like neutrophils, and abundant infiltration of anti-tumorigenic macrophages in the pancreatic preneoplastic microenvironment. Mechanistically, we demonstrate that nSMase2-expressing PDA cell small extracellular vesicles (sEVs) reduce survival of KPC mice; PDA cell sEVs generated independently of nSMase2 prolong survival of KPC mice and reprogram macrophages to a proinflammatory phenotype. Collectively, our study highlights previously unappreciated opposing roles for exosomes, based on biogenesis pathway, during PDA progression.

**Graphical abstract:** 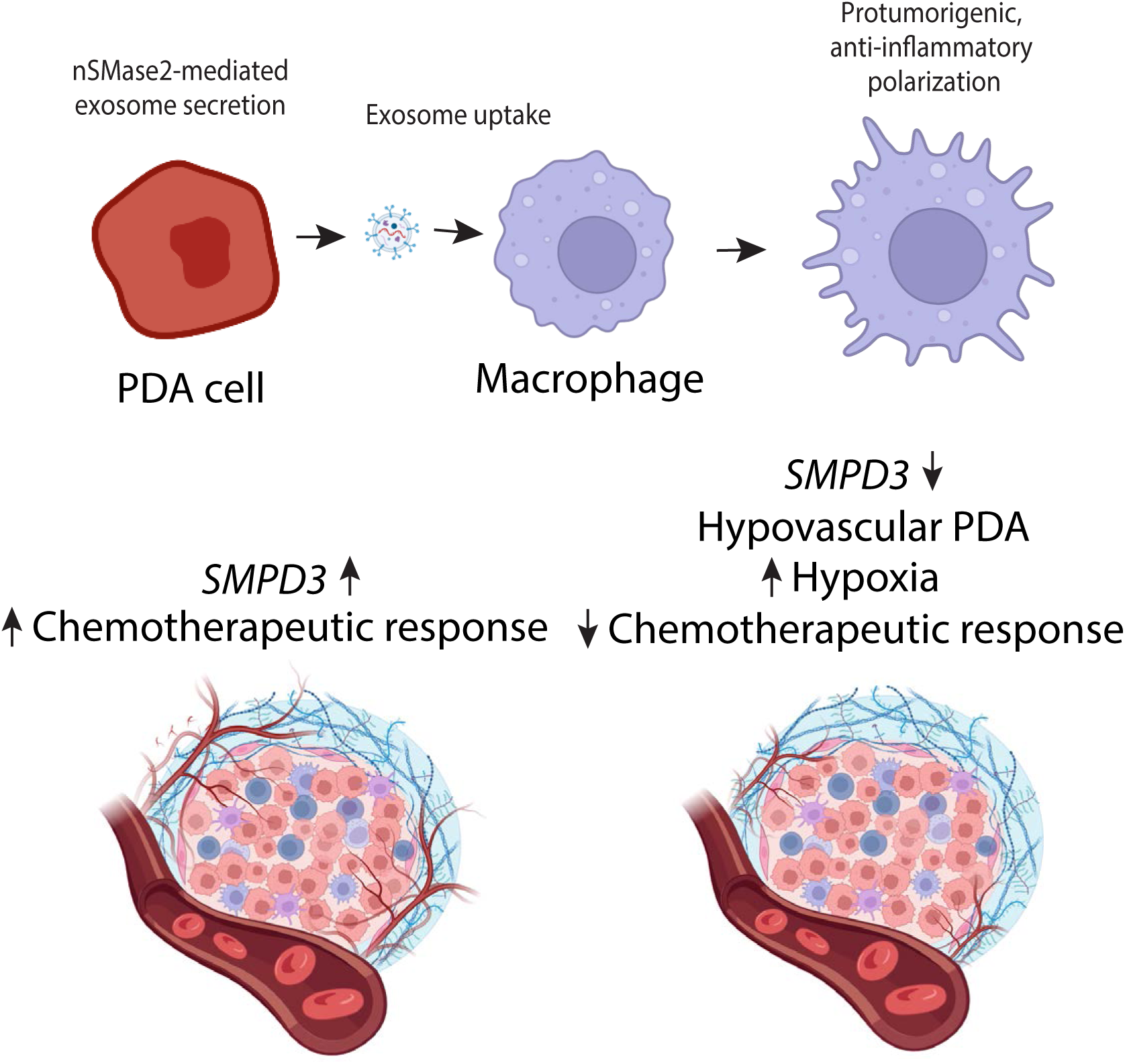

## INTRODUCTION

Exosomes are membrane-enclosed, small extracellular vesicles (sEVs) of endocytic origin ranging in size from ∼40nm to 160nm that are secreted by all cells during both normal physiology and abnormal cellular states. A variety of bioactive molecules are packaged into exosomes including nucleic acids (small RNAs, mRNA, lncRNA, DNA), proteins, metabolites, and lipids (Kalluri and LeBleu, 2020). Exosomes have been hypothesized to be important mediators of intercellular communication as their transfer of cargo following uptake by recipient cells has demonstrated functional relevance (Valadi et al., 2007). Exosome biogenesis can occur through several mechanisms including a ceramide-dependent pathway that promotes the initial inward budding of endosomal membranes to form intraluminal vesicles (Trajkovic et al., 2008), a pathway that is independent of endosomal sorting complexes required for transport (ESCRT) (Schmidt and Teis, 2012).

Exosomes produced by pancreatic ductal adenocarcinoma (PDA) cells can functionally influence tumorigenesis and serve as biomarkers for disease progression. Costa-Silva et al. demonstrated that PDA cell exosomes initiate pre-metastatic niche formation in the liver (Costa-Silva et al., 2015). Pancreatic cancer cell derived sEVs carry double stranded genomic DNA with mutated KRAS and p53 (Kahlert et al., 2014) in addition to expressing the cell surface proteoglycan Glypican-1, a biomarker that distinguishes healthy patients from PDA patients (Melo et al., 2015). A number of studies have implicated alteration in function and expression of exosome biogenesis genes in human cancer including PDA (Albacete-Albacete et al., 2020; Gheytanchi et al., 2021; Wei et al., 2015; Wu et al., 2023; Zhao et al., 2016). Exosome biogenesis pathways influence packaging of exosome constituents. The neutral sphingomyelinase 2 (nSMase2), which is encoded by sphingomyelin phosphodiesterase 3 (*Smpd3*), catalyzes the hydrolysis of sphingomyelin to ceramide and phosphocholine; ceramide generated by nSMase2 is thought to be a major source for the ceramide-dependent exosome biogenesis pathway. The biological significance of nSMase2-mediated exosome secretion during PDA development remains unclear.

Here, we explored the intrinsic and extrinsic roles of nSMase2 on PDA cell biology and exosome secretion. Regarding intrinsic function, nSMase2 loss alters the lipid profile and gene expression of PDA cells and promotes a subtype switch from basal to classical. Concerning extrinsic, we investigated how PDA cell exosome cargo heterogeneity and function are impacted by the nSMase2-mediated exosome biogenesis pathway. We unveil that PDA cell sEVs generated through the ceramide-dependent pathway are protumorgenic during PDA progression: they thwart survival of KPC (*Ptf1a-Cre^Tg/wt^*; *Kras^Tg/wt^*; *Trp53^f/wt^*) mice, a mouse model of PDA (von Figura et al., 2014). PDA cell sEVs generated independently of nSMase2 are antitumorigenic: they polarize macrophages toward a proinflammatory phenotype and prolong survival of KPC mice. Our findings demonstrate a role for PDA cell nSMase2-generated exosomes in driving PDA.

## RESULTS

### sEV secretion by pancreatic cells is variably affected by neutral sphingomyelinase activity

We detected nSMase2 expression and nSMase enzymatic activity in a panel of murine and human PDA cell lines as well as the immortalized human pancreatic duct cell lines hTERT-HPNE and HPDE6c7 (Figure 1 A-C, Supplemental Figure 1A). Polyclonal murine PDA cell lines were generated from end-stage KPC (*Ptf1a-Cre^Tg/wt^*; *Kras^Tg/wt^*; *Trp53^f/wt^*) (von Figura et al., 2014) mouse pancreatic tumors. To assess the ability of nSMase2 to regulate pancreatic cell sEV secretion, we next introduced a CD63eGFP construct (murine CD63 cDNA fused to eGFP) into murine PDA cell lines and the pCT-CD63-GFP (human CD63 fused to GFP) construct into human pancreatic cell lines through viral transduction and selection for stable expression to allow for quantification of CD63^+^ sEV secretion. The tetraspanin CD63 has been proposed to be highly enriched in exosomes (Escola et al., 1998; Kowal et al., 2016). Then, we treated a subset of these CD63eGFP -expressing murine and CD63-GFP-expressing human pancreatic cell lines with the nSMase inhibitor GW4869 (Risner et al., 2023; Tabatadze et al., 2010). Equal numbers of cells were seeded, treated with 20uM GW4869 or vehicle control for 24 hours, and CD63^+^ sEVs isolated from conditioned media were subsequently quantified using NanoSight and normalized to cell number. We observed that GW4869 treatment significantly reduced CD63^+^ sEV secretion from hTERT-HPNE CD63-GFP and MIA PaCa-2 CD63-GFP cell lines and did not affect CD63^+^ sEV secretion from PANC-1 CD63-GFP, CFPAC-1 CD63-GFP, *KPC; Smpd3^wt/wt^* 7 CD63eGFP, or *KPC; Smpd3^wt/wt^* 14 CD63eGFP cell lines (Figure 1D). These data suggest that nSMase2 modulation of sEV secretion from pancreatic cells is context dependent, an observation in line with previous reports (Choezom and Gross, 2022; Kosaka et al., 2010; Phuyal et al., 2014; Poggio et al., 2019).

**Figure 1.**
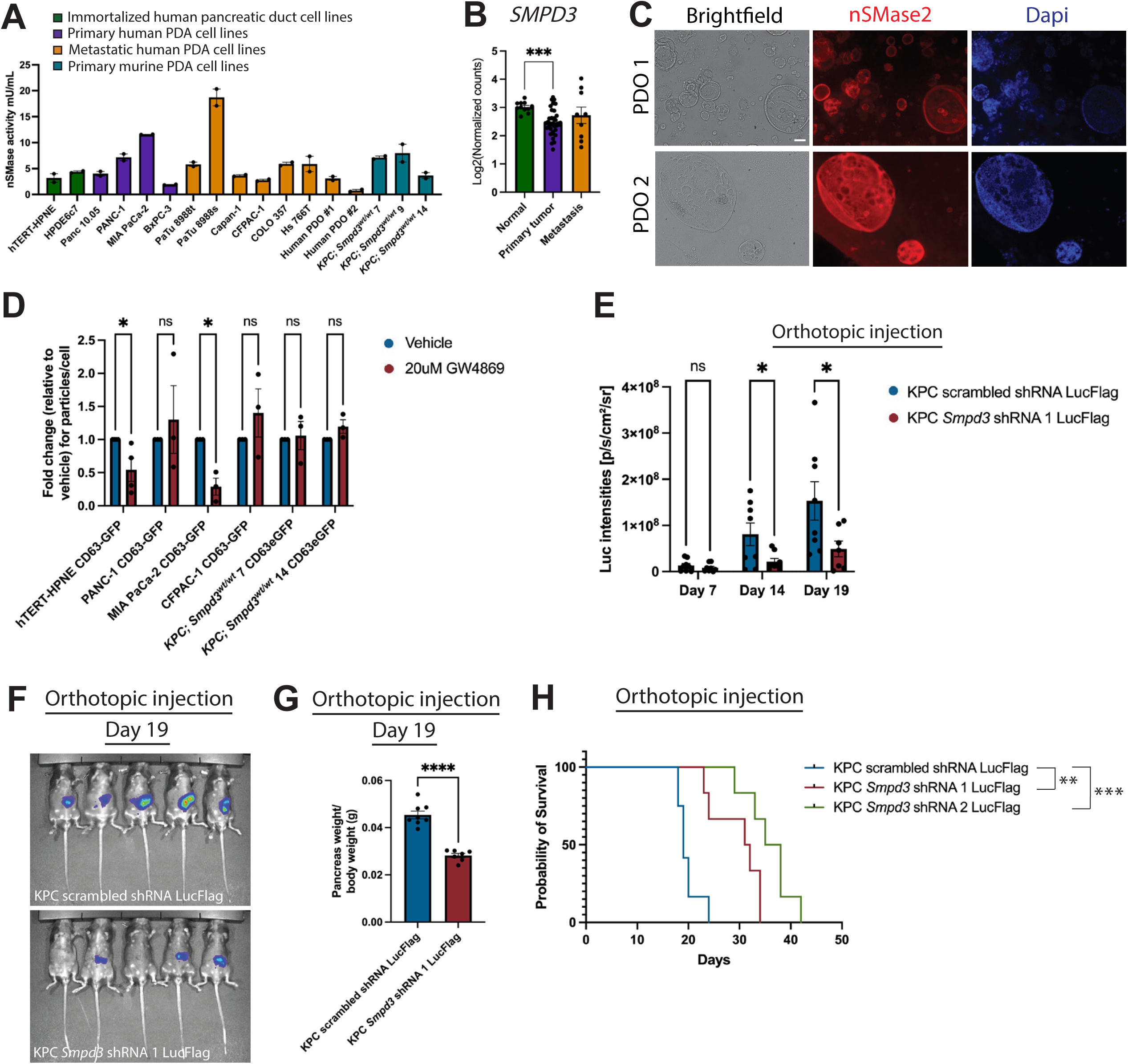
*Smpd3* modulates pancreatic tumor growth. A) Immortalized human pancreatic duct cells (hTERT-HPNE and HPDE6c7), human and murine PDA cell lines, and human patient derived-organoids (PDOs) demonstrate nSMase activity. Values shown are technical replicates of one lysate per cell line. B) *SMPD3* expression was analyzed in an existing human pancreas organoid RNA-Seq dataset (Tiriac et al., 2018) and is depicted in normal (n=11), primary (n=35), and metastatic (n=9) human organoids. Log of normalized counts of *SMPD3* expression are plotted. C) Immunocytochemistry demonstrates nSMase2 expression in two pancreatic metastatic PDOs. Scale bar is 100uM. D) Nanoparticle tracking analysis of CD63^+^ sEVs upon GW4869 and vehicle treatment is depicted. E) IVIS imaging results at Day 7, 14, and 19 are graphed. Eight animals were injected with each cell line. F) Images depict bioluminescence signal during IVIS imaging at day 19. G) Pancreas weight/body weight at day 19 is graphed. Average values for the KPC scrambled shRNA are 0.04540 ± 0.001635 (n=8) and KPC *Smpd3* shRNA 1 are 0.02817 ± 0.0008858 (n=7). H) Rag 1 KO mice injected orthotopically with KPC *Smpd3* shRNA1 and KPC *Smpd3* shRNA2 cell lines lived significantly longer than Rag1 KO mice injected orthotopically with KPC scrambled shRNA cells (*P*=0.0013 using log-rank test and *P*=0.0001 using log-rank test, respectively). The median survival of Rag 1 KO mice injected orthotopically with KPC scrambled shRNA cells (n=12) is 19 days, KPC *Smpd3* shRNA 1 cells (n=6) is 31.50 days, and KPC *Smpd3* shRNA 2 cells (n=6) is 36.50 days.

To assess the ability of nSMase2 to regulate pancreatic cancer cell growth, we genetically modified the polyclonal primary cell lines from KPC mouse pancreatic tumors to stably express shRNAs targeting *Smpd3* and a flag-tagged firefly luciferase construct to allow for *in vivo* tumor cell growth monitoring (Supplemental Figure 1B). Next, we injected *Smpd3* knockdown KPC and scrambled shRNA control KPC cells both subcutaneously and orthotopically into Rag1 KO mice. Subcutaneous injection of the KPC *Smpd3* shRNA 2 cell line resulted in modestly reduced tumor volume at day 17 when compared to the KPC scrambled shRNA cell line (Supplemental Figure 1C). At day 17 post injection, the weight of tumors from both KPC *Smpd3* shRNA 1 and KPC *Smpd3* shRNA 2 cell lines was significantly less than the KPC scrambled shRNA cell line (Supplemental Figure 1D). Orthotopic injection of the *Smpd3* knockdown KPC and control KPC cells that express a flag-tagged firefly luciferase demonstrated that *Smpd3* knockdown significantly reduces pancreatic tumor growth *in vivo* as early as 14 days after injection (Figure 1E-F) as well as pancreas weight/body weight ratio (Figure 1G) and extends survival of Rag1 KO mice (Figure 1H) when compared to the KPC scrambled shRNA control. Together, these data suggest that *Smpd3* regulates pancreatic tumor growth *in vivo*.

### *Smpd3* knockdown in PDA cells reduces proliferation of epithelial PDA cells *in vivo*

nSMase2 has been shown to regulate the cell cycle (Marchesini et al., 2004; Maupas-Schwalm et al., 2009). Given this, we next assessed proliferation in tumors generated from the orthotopically injected *Smpd3* knockdown KPC and control KPC cells. Tumors generated using the KPC *Smpd3* shRNA 2 cell line had significantly fewer Ki67^+^ epithelial cells than tumors generated using the KPC scrambled shRNA cell line (Supplemental Figure 1E). Various cellular stresses including oxidative (Levy et al., 2006) and chemotherapy-induced (Ito et al., 2009) have been shown to upregulate nSMase2 expression and induce apoptosis. Thus, we next queried whether levels of epithelial *Smpd3* expression affected apoptosis in the *in vivo* injected tumors. We observed no difference in cleaved caspase 3 positive epithelial cells in tumors generated from the orthotopically injected *Smpd3* knockdown KPC and control KPC cells (Supplemental Figure 1F).

Using cell viability (CTG) and colony formation assays, the *Smpd3* knockdown KPC cell lines did not display a difference in metabolic activity or proliferation *in vitro* (Supplemental Figure 1G); yet *Smpd3* knockdown KPC cells proliferated less *in vivo* when injected into Rag1 KO mice, suggesting that nSMase2 in conjunction with the tumor microenvironment (TME) likely influenced the growth and development of the pancreatic tumors. Additionally, we noted that the impact on pancreas tumor volume and weight in the subcutaneous tumors was subtler than that observed in the orthotopic model, suggesting that the pancreatic niche is important for nSMase2-mediated modulation of pancreatic tumor growth. A major component of the TME that influences PDA phenotype is the extracellular matrix (ECM) stroma. Thus, to examine if reduction of epithelial nSMase2 in KPC cells affected the ECM, we next quantified fibrillar collagen in tumors generated from the cell lines using picrosirius red staining and polarized light imaging to visualize. Analysis revealed a modest decrease in fibrillar collagens in the tumor tissues generated using the KPC *Smpd3* shRNA 2 cell line when compared to the KPC scrambled shRNA cell line (Supplemental Figure 1H-I). Summarily, these data suggest that reducing PDA cell *Smpd3* expression decreases proliferation of PDA cells *in vivo*.

### Pancreatic loss of epithelial nSMase2 delays neoplasia formation and extends survival of KPC mice

The subcutaneous and orthotopic studies suggested *Smpd3* influences PDA tumor phenotype and implicated the TME. Nevertheless, given the limitation of cell line studies to assess the TME phenotype, we next generated genetically engineered mouse models (GEMMs) of PDA in which *Smpd3* was knocked out. We generated an *Smpd3* floxed mouse (Supplemental Figure 2A) and crossed these mice to the KPC (*Ptf1a-Cre^Tg/wt^*; *Kras^Tg/wt^*; *Trp53^f/wt^*) mouse model of pancreatic cancer. *Smpd3* is expressed in pancreatic preneoplasia including acinar-to-ductal-metaplasia (ADM) and pancreatic intraepithelial neoplasia (PanIN) and neoplasia in KPC mice (Supplemental Figure 2B-D). *Ptf1a-Cre^Tg/wt^; Smpd3^f/f^*mice were born in Mendelian ratios and displayed no histological phenotype at 1 year of age (Supplemental Figure 2E). Survival studies revealed that the *KPC; Smpd3^f/f^* mice lived significantly longer than *KPC; Smpd3^wt/wt^* mice (Figure 2A). Loss of epithelial nSMase2 in KPC mice did not affect tumor histological classification or occurrence of macrometastasis at end-stage (Figure 2B-C; Supplemental Figure 2F). However, *KPC; Smpd3^f/f^* mice had significantly fewer PanIN lesions than *KPC; Smpd3^wt/wt^* mice, as measured by alcian blue staining, at 10-11 weeks of age (Figure 2D-E). There was also a modest decrease in edema in the *KPC; Smpd3^f/f^* mice when compared to *KPC; Smpd3^f/wt^* mice at 10-11 weeks of age (Figure 2D,F). Quantification of tumor area using hematoxylin and eosin-stained sections of 19–21-week-old mice harboring PDA showed a significant decrease in tumor burden in *KPC; Smpd3^f/f^* mice when compared to *KPC; Smpd3^wt/wt^*mice (Figure 2G-H). Together, these data suggest that in cooperation with oncogenic *Kras* and heterozygous *Trp53* loss, epithelial nSMase2 expression is protumorigenic during PDA development and progression.

**Figure 2.**
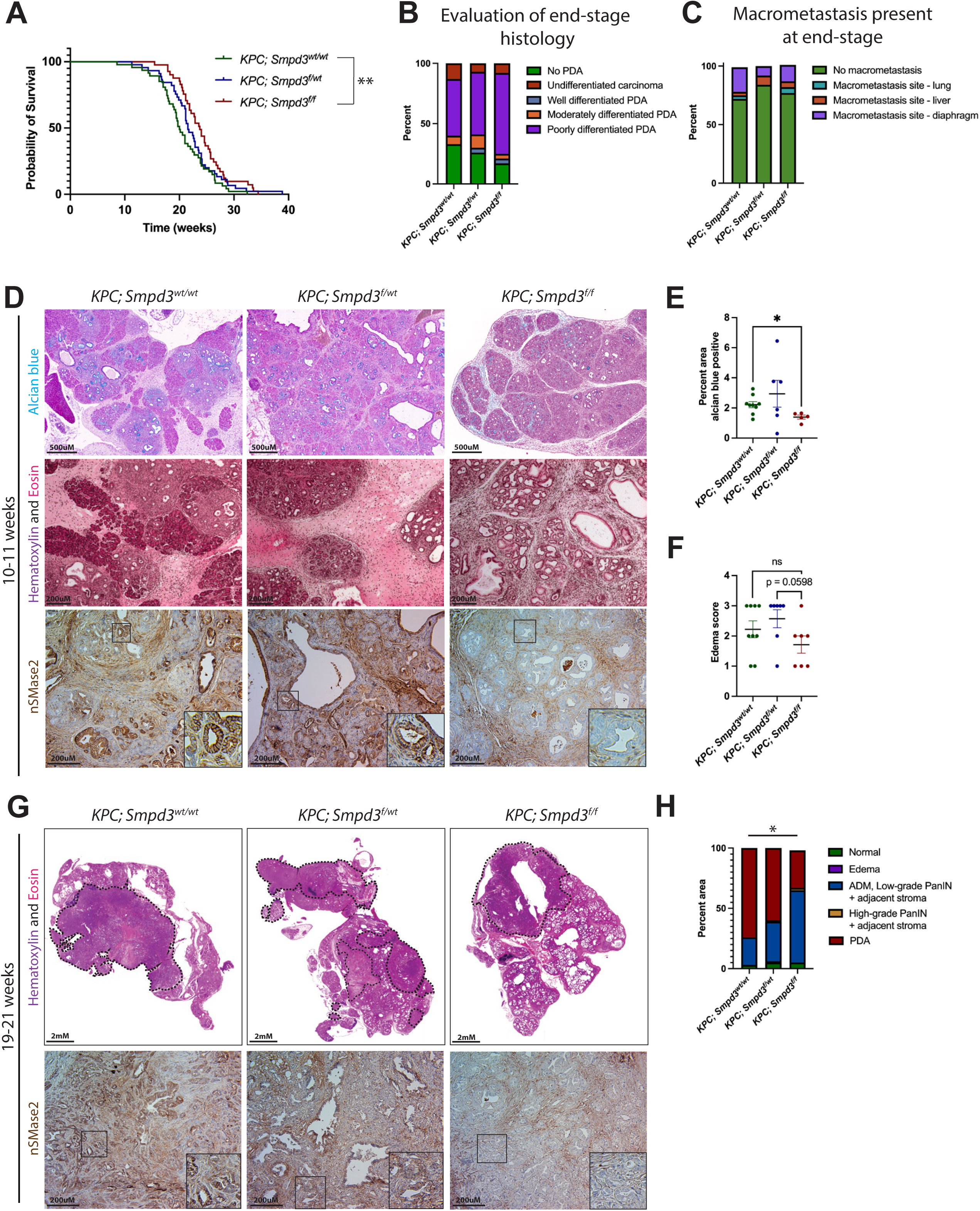
*Smpd3* ablation reduces formation of neoplasia and prolongs survival of KPC mice. A) *KPC; Smpd3^f/f^* mice lived significantly longer than *KPC; Smpd3^wt/wt^*mice (*P*=0.0039 using log-rank (Mantel-Cox) test). The median survival of *KPC; Smpd3^f/f^* mice (n=41) is 23.71 weeks, *KPC; Smpd3^f/wt^* mice is 21.57 weeks (n=45), and *KPC; Smpd3^wt/wt^* mice (n=47) is 20.00 weeks. B) The percentage of animals having PDA at end-stage in addition to the differentiation status of primary pancreatic tumors at end-stage is graphed for *KPC; Smpd3^wt/wt^* mice (n=30), *KPC; Smpd3^f/wt^* mice (n=27), and *KPC; Smpd3^f/f^* mice (n=24). (C) The percentage of animals having macrometastasis at end-stage in addition to the site of macrometastasis at end-stage is depicted for *KPC; Smpd3^wt/wt^* (n=29), *KPC; Smpd3^f/wt^* (n=25), and *KPC; Smpd3^f/f^* (n=22) mice. Using Fisher’s exact tests, no significant differences were observed in the number or site of macrometastatic lesions present at end-stage in *KPC; Smpd3^wt/wt^*, *KPC; Smpd3^f/wt^*, or *KPC; Smpd3^f/f^* mice. If macrometastasis was present, only one site was observed per animal in these cohorts. D-E) Quantification of alcian blue staining demonstrates significantly reduced PanIN lesions in *KPC; Smpd3^f/f^* mice (n=5) when compared to *KPC; Smpd3^wt/wt^* mice (n=9) (*P*=0.0154). H&Es used to score edema are depicted. nSMase2 immunohistochemistry depicts loss of nSMase2 in pancreatic epithelium of PanIN-bearing *KPC; Smpd3^f/f^*mice. F) Edema scoring for *KPC; Smpd3^wt/wt^* (n=9), *KPC; Smpd3^f/wt^* (n=7), and *KPC; Smpd3^f/f^*(n=7) mice is depicted. The scoring system used is 1=low, 2=medium, and 3=high. G-H) The percentage of pancreatic area occupied by PDA is significantly higher in *KPC; Smpd3^wt/wt^* mice (74.16 ± 8.705) when compared to *KPC; Smpd3^f/f^* mice (30.51 ± 17.72) (*P*=0.0497) at 19-21 weeks of age. Average percent area is graphed in H (n=4 *KPC; Smpd3^f/f^*mice, n=4 *KPC; Smpd3^f/wt^* mice, n=5 *KPC; Smpd3^wt/wt^*mice). Dotted lines show the PDA area in representative images. nSMase2 immunohistochemistry depicts loss of nSMase2 in pancreatic epithelium of PDA-bearing *KPC; Smpd3^f/f^* mice at 19-21 weeks of age.

### nSMase2 does not alter KPC; Smpd3^wt/wt^, KPC; Smpd3^f/wt^, and KPC; Smpd3^f/f^ tumor cell growth in vitro

To comprehensively interrogate the intrinsic functions of *Smpd3*, including how altering expression levels affects pancreatic cancer cells on a molecular level, we next generated multiple polyclonal primary cell lines from end-stage primary and metastatic pancreatic tumors of *KPC; Smpd3^wt/wt^* (n=19), *KPC; Smpd3^f/wt^*(n=14), and *KPC; Smpd3^f/f^* (n=17) mice (Supplemental Figure 3A-E, G, Supplemental Table 1). Because mutations which reduce or eliminate *SMPD3* gene expression include deep deletions and missense mutations as well as alterations that increase *SMPD3* gene expression, such as genomic amplifications, have been reported in PDA patients (Consortium, 2020), we generated KPC cell lines that overexpress *Smpd3* as well (Supplemental Figure 3F-G). To determine if nSMase2 levels affect proliferation *in vitro* in the polyclonal murine PDA cell lines, we performed two cell growth assays. For the first assay, equal numbers of cells were seeded in media containing exosome-depleted FBS, and cell number was estimated 6 days later using the Cell Titer Glow (CTG) viability assay, an assay that quantifies viable, metabolically active cells by estimating cellular ATP levels. Similar to the *Smpd3* knockdown KPC cell lines, we observed no difference in metabolically active cells in the polyclonal murine tumor cell lines generated from *KPC; Smpd3^wt/wt^*, *KPC; Smpd3^f/wt^*, and *KPC; Smpd3^f/f^* pancreata (Figure 3A). For the second growth assay, we seeded 100 cells in media containing exosome-depleted FBS, and quantified colony formation after 11 days. No differences in colony formation were observed (Figure 3B). These data suggest that nSMase2 does not alter pancreatic cancer epithelial cell proliferation *in vitro*; once again suggesting that nSMase2 mediated exosome secretion in conjunction with the TME, likely plays a role in development and progression of PDA.

**Figure 3.**
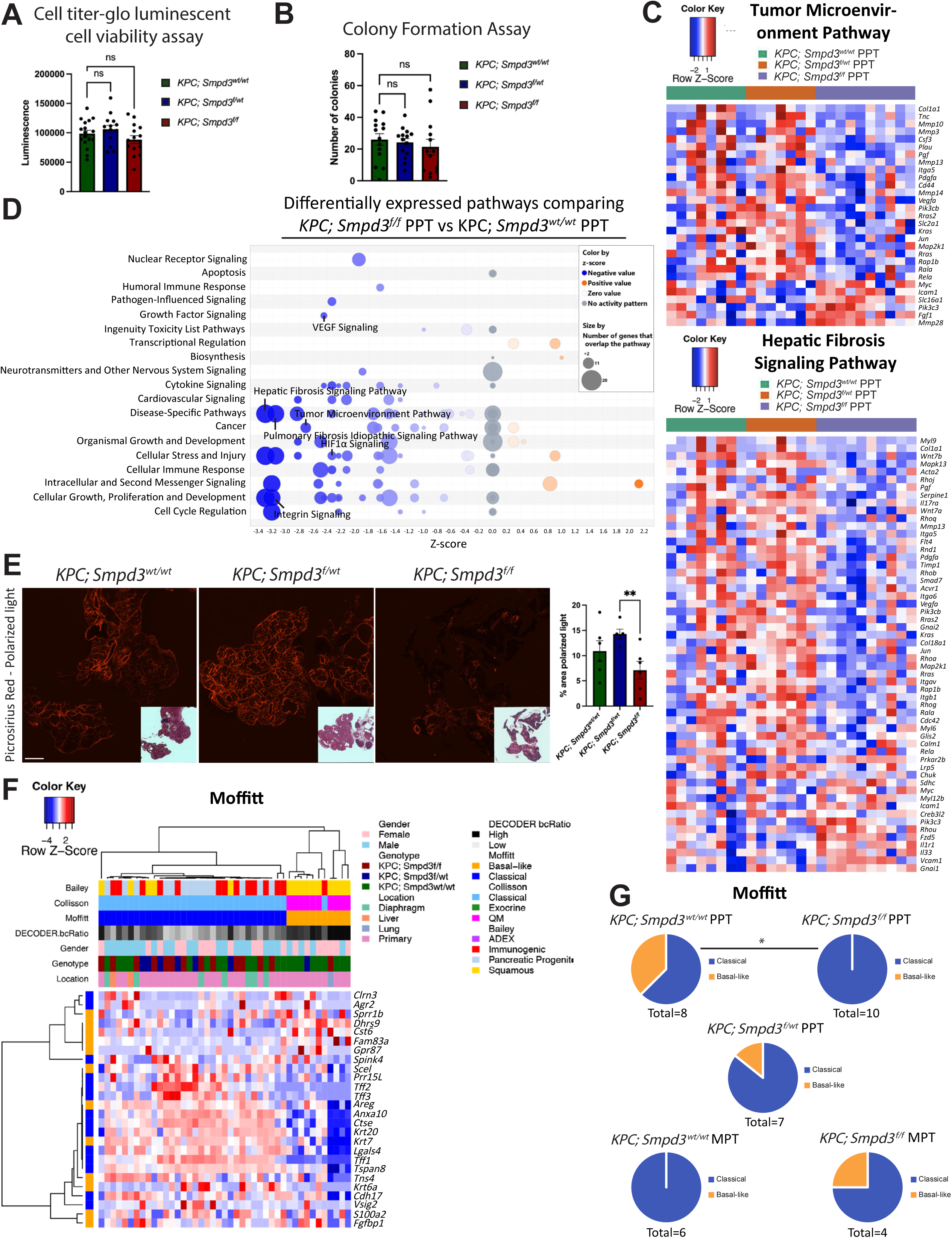
RNA sequencing demonstrates a role for *Smpd3* in modulation of cellular pathways and regulation of PDA subtype. A) Average luminescence values from 3-5 independent experiments per cell line are graphed. Average values for primary PDA cell lines generated from *KPC; Smpd3^wt/wt^* mice (n=17 cell lines) are 98,614 ± 5,684, *KPC; Smpd3^f/wt^* mice (n=14 cell lines) are 106,061 ± 6,699, and *KPC; Smpd3^f/f^* mice (n=15 cell lines) are 88,460 ± 6,672. B) Average number of colonies from at least 3 independent colony formation assays per cell line are depicted. Average values from *KPC; Smpd3^wt/wt^* mice (n=14 cell lines) are 26.02 ± 3.652, *KPC; Smpd3^f/wt^* mice (n=14 cell lines) are 24.16 ± 2.519, and *KPC; Smpd3^f/f^* mice (n=13 cell lines) are 21.34 ± 4.872. C) Heat maps depict expression of significantly deregulated molecules in Tumor Microenvironment Pathway and Hepatic Fibrosis Signaling Pathway in all three PPT cell lines. Molecules shown in heat maps include those molecules within the Ingenuity pathway analysis (IPA) pathway that were significantly deregulated in any comparison between PPT groups. D) IPA results show a bubble category chart of significantly deregulated pathways when comparing *KPC; Smpd3^f/f^* PPT and *KPC; Smpd3^wt/wt^* PPT cell lines. A positive z-score represents upregulation, and a negative z-score indicates downregulation of a pathway in *KPC; Smpd3^f/f^* PPT when compared to *KPC; Smpd3^wt/wt^* PPT cell lines. A gray circle depicts significant overrepresentation of a pathway, the direction of which cannot yet be determined. Select pathways are annotated. E) Picrosirius red stained images are shown for *KPC; Smpd3^wt/wt^* (n=6), *KPC; Smpd3^f/wt^* (n=5), and *KPC; Smpd3^f/f^* (n=6) mice at 10-11 weeks of age. The small piece of intestine in the lower left corner of the *KPC; Smpd3^f/wt^* image was excluded from the quantification. Scale bar is 1000uM. Inset shows brightfield image. F) Transcriptional subtyping of end-stage pancreatic tumor cell lines generated from *KPC; Smpd3^wt/wt^*(PPT and MPT), *KPC; Smpd3^f/wt^* (PPT), *KPC; Smpd3^f/f^*(PPT and MPT) as well as *KPC* Mock OE and *KPC Smpd3* OE lines is depicted. Genes used to identify subtypes according to the Moffitt classification are shown. G) Pie charts depict the number of cell lines with the classical and basal-like Moffitt subtypes. When comparing *KPC; Smpd3^wt/wt^* PPT vs *KPC; Smpd3^f/f^*PPT using Moffitt classification and Chi square test, p-value=0.0339.

### *Smpd3* regulates intracellular pathways governing TME dynamics in PDA cells

Intracellular nSMase2-generated ceramide signaling has been shown to function as a second messenger molecule in cellular stress responses (Clarke and Hannun, 2006). To examine the intrinsic effects of *Smpd3* loss in PDA cells, we performed RNA-sequencing of the polyclonal murine PDA cell lines, including cell lines generated from primaries and metastases and the overexpression cell lines (Supplemental Table 1). Differentially expressed gene (DEG) lists (Supplemental Table 2) were used in ingenuity pathway analysis (IPA). When comparing *KPC; Smpd3^f/f^* primary pancreatic tumor (PPT) cell lines vs *KPC; Smpd3^wt/wt^* PPT cell lines, multiple pathways that modulate the TME including Tumor Microenvironment Pathway and Hepatic Fibrosis Signaling Pathway were significantly downregulated in *KPC; Smpd3^f/f^* PPT cell lines (Figure 3C-D and Supplemental Table 2). Pathways regulating cell adhesion and mechanotransduction including Integrin Signaling, Epithelial Adherens Junction Signaling, and Actin Cytoskeleton Signaling were also significantly downregulated in *KPC; Smpd3^f/f^* PPT cell lines when comparing *KPC; Smpd3^f/f^* PPT vs *KPC; Smpd3^wt/wt^* PPT cell lines (Figure 3D, Supplemental Table 2, and Supplemental Figure 4A). miR-802, which suppresses pancreatic cancer initiation (Ge et al., 2022), was predicted to be activated in *KPC; Smpd3^f/f^* PPT by IPA Upstream Regulator Analysis when comparing *KPC; Smpd3^f/f^* PPT to *KPC; Smpd3^wt/wt^* PPT (Supplemental Table 2). Additionally, mediators of inflammation and immune response such as NFkB (complex), IL1 and CD40, were predicted to be activated in *KPC; Smpd3^f/wt^* PPT by IPA Upstream Regulator Analysis when comparing *KPC; Smpd3^f/wt^* PPT to *KPC; Smpd3^wt/wt^* PPT (Supplemental Table 2).

Next, we assessed transcript expression differences comparing all three PPT groups (*KPC; Smpd3^wt/wt^*, *KPC; Smpd3^f/wt^*, and *KPC; Smpd3^f/f^*polyclonal PPT cell lines) utilizing Sleuth (Pimentel et al., 2017). An observation consistent with the delayed neoplasia formation, reduced tumor burden, and extended survival in *KPC; Smpd3^f/f^* mice, multiple genes previously reported to promote PDA progression in human models were identified as upregulated in *KPC; Smpd3^wt/wt^*cell lines and downregulated in *KPC; Smpd3^f/f^* cell lines including *Ano1* (Zhang et al., 2023), *Actn1* (Rajamani and Bhasin, 2016), *Vegfa* (Sun et al., 2007), *Ilk* (Sawai et al., 2006), *Fn1* (Hu et al., 2019), *Cd151* (Zhu et al., 2011), *Ccn2* (Bennewith et al., 2009), *Anxa1* (Pessolano et al., 2018), *Pvr* (Nishiwada et al., 2015), *Ly6e* (Kihara, 2016), *Hmga1* (Chia et al., 2023), *Msn* (Torer et al., 2007), and *Serpine1* (Liu et al., 2018) (Supplemental Figure 4B and Supplemental Table 2).

When comparing *KPC; Smpd3^f/f^* metastatic pancreatic tumor (MPT) cell lines vs *KPC; Smpd3^wt/wt^* MPT cell lines, multiple pathways regulating metabolism including Pentose Phosphate Pathway, Purine Nucleotides De Novo Biosynthesis II, and Production of Nitric Oxide and Reactive Oxygen Species in Macrophages were enriched in *KPC; Smpd3^f/f^*MPT cells (Supplemental Figure 4C-D and Supplemental Table 2). When comparing *KPC Smpd3* OE vs *KPC* Mock OE, only six pathways were significantly differentially regulated, which may be due in part to the modest level of *Smpd3* expression increase in overexpression lines (Supplemental Figure 4E-F and Supplemental Table 2). Collectively, the RNA-Seq analyses show that nSMase2 modulates PDA cell gene expression.

Given the consistent observation of alteration of expression of genes involved in fibrosis in the transcriptomic analyses, we next assessed protein expression of fibrillar collagens, which are the main ECM component in the pancreas stroma, using picrosirius red staining followed by polarized light imaging and abundance of two predominant ECM-secreting pancreatic cell types – fibroblasts and stellate cells. We observed protein expression of collagen was significantly reduced in *KPC; Smpd3^f/f^* pancreata when compared to *KPC; Smpd3^f/wt^* pancreata (Figure 3E). Quantification of activated pancreatic stellate cells (PSCs) defined by αSMA^+^GFAP^+^Desmin^+^Dapi^+^ copositive and activated fibroblasts defined by αSMA^+^Vimentin^+^FAP^+^Dapi^+^ copositive in *KPC; Smpd3^wt/wt^*, *KPC; Smpd3^f/wt^*; and *KPC; Smpd3^f/f^* pancreata at the 10-11 week time point revealed *KPC; Smpd3^f/f^* pancreata display significantly fewer activated PSCs and fibroblasts when compared to both *KPC; Smpd3^f/wt^* and *KPC; Smpd3^wt/wt^* pancreata (Supplemental Figure 5A-D). These *in vivo* observations validate a subset of the RNA-seq analyses on PDA cells.

### *Smpd3* loss results in PDA subtype switching from aggressive basal-like to classical

PDA subtypes have been described and can be determined using bulk RNA-sequencing data (Bailey et al., 2016; Collisson et al., 2019; Moffitt et al., 2015; Peng et al., 2019). We next queried whether the murine polyclonal PDA cell lines generated from end-stage pancreatic tumors formed different subtypes of PDA (Figure 3F and Supplemental Table 2). Using 50 tumor specific genes from Moffitt et al.’s study for subtyping analysis, we observed a shift in PDA subtype from the more aggressive basal-like subtype seen in 3/8 *KPC; Smpd3^wt/wt^* PPTs to classical, seen in 10/10 *KPC; Smpd3^f/f^* PPTs (Figure 3F-G). *Smpd3* overexpression caused a shift from basal to classical in two *KPC; Smpd3^wt/wt^* PPT cell lines using the Moffitt classification (Figure 3F, Supplemental Figure 5E, and Supplemental Table 2). PPT and MPT cell lines that were identified as classical in the Moffitt scheme were also identified as classical in the Collisson scheme, and PPT and MPT cell lines that were identified as basal-like in the Moffitt scheme identified as quasi-mesenchymal in the Collisson classification (Figure 3F and Supplemental Table 2). The Exocrine-like subtype in the Collisson scheme and the ADEX subtype in the Bailey scheme were not identified in the dataset. Consistent with phenotypic observations in the mouse models, these data suggest that *Smpd3* loss in epithelial pancreatic cells cooperates with oncogenic *Kras* and heterozygous *Trp53* loss to promote formation of a classical PDA subtype (Collisson et al., 2019).

### *Smpd3* directs biogenesis of ultra-long chain ceramide species in PDA cells

nSMase2 generates ceramide and phosphocholine from sphingomyelin. IPA of the RNA-seq data showed that Ceramide Signaling is significantly decreased in *KPC; Smpd3^f/f^* PPT when compared to *KPC; Smpd3^f/wt^*PPT (Supplemental Figure 6A and Supplemental Table 2). Thus, we next analyzed the lipid composition of the polyclonal murine PDA cell lines using mass spectrometry (Supplemental Table 3). The lipid profile of the polyclonal murine PDA cell lines contained significantly differentially expressed fatty acyls, glycerolipids, glycerophospholipids, organooxygen compounds, and sphingolipids (Supplemental Figure 6B-C). When comparing *KPC; Smpd3^f/f^*PPT to *KPC; Smpd3^wt/wt^* PPT, a significant reduction in ultra-long chain (Horbay et al., 2022) ceramides including d32:1, d34:0, d34:1, d34:2, and d36:1 was observed (Supplemental Figure 6D and Supplemental Table 3). Comparison of *KPC Smpd3* OE to *KPC* Mock OE cell lines showed no significantly differentially expressed ceramide species; however, two phosphatidylcholine (phosphocholine is an intermediate in phosphatidylcholine synthesis) species – 42:8 and O-34:4 – are significantly upregulated in *KPC Smpd3* OE (Supplemental Figure 6E and Supplemental Table 3). Our data show that nSMase2 is a critical regulator of the lipid profile of PDA cells.

### nSMase2-generated PDA cell exosomes thwart survival of KPC mice

As demonstrated by the CD63^+^ sEV quantification (Supplemental Figure 7A), *KPC; Smpd3^f/f^* PDA cells still secrete CD63^+^ sEVs in the absence of nSMase2, which is unsurprising given the presence of alternate mechanisms for sEV biogenesis within a cell (McAndrews and Kalluri, 2019). However, nSMase2 has been shown to affect the cellular constituents packaged into exosomes (Guo et al., 2015; Kosaka et al., 2010). To test the hypothesis that nSMase2-generated exosome content affects PDA progression, we next administered equal numbers of purified sEVs isolated from the murine polyclonal PDA PPT cell lines using the Cushioned-Density Gradient Ultracentrifugation (C-DGUC) protocol (Duong et al., 2019) (Supplemental Figure 7B) intraperitoneally (IP) thrice weekly to *KPC; Smpd3^wt/wt^* and *KPC; Smpd3^f/f^* mice beginning at 6 weeks of age and assessed survival (Figure 4A). Kamerkar et al. previously showed that IP injection of a conservative dose of 10^8^ sEVs every other day into mice resulted in injected sEV uptake by pancreatic cells (Kamerkar et al., 2017). *KPC; Smpd3^wt/wt^*mice injected with 1 x 10^8^ sEVs thrice weekly isolated from *KPC; Smpd3^f/f^* PDA PPT cell lines (n=17) survived significantly longer than the control - *KPC; Smpd3^wt/wt^* mice injected with 1 x 10^8^ sEVs thrice weekly isolated from *KPC; Smpd3^wt/wt^* PDA PPT cell lines (n=17), suggesting that sEVs secreted from *KPC; Smpd3^f/f^* PDA PPT cells are anti-tumorigenic (Figure 4B). *KPC; Smpd3^f/f^* mice injected with sEVs isolated from *KPC; Smpd3^wt/wt^* PDA PPT cell lines (n=11) displayed significantly shortened life spans when compared to the respective control, *KPC; Smpd3^f/f^* mice injected with sEVs isolated from *KPC; Smpd3^f/f^* PDA PPT cell lines (n=13), suggesting that *KPC; Smpd3^wt/wt^* PDA PPT cell sEVs are protumorigenic during PDA progression (Figure 4B). Using Fisher’s exact tests, we observed no difference in the incidence of primary tumors, primary tumor differentiation, incidence of macrometastasis, or the site of macrometastasis in sEV injected models (Supplemental Figure 7C-D). However, injecting mice with sEVs resulted in a significant change in pancreas weight/body weight ratio when comparing *KPC; Smpd3^wt/wt^* uninjected to *KPC; Smpd3^wt/wt^*injected with sEVs isolated from *KPC; Smpd3^f/f^* PDA PPT cell lines (Supplemental Figure 7E-F). We also observed a prevalence of pancreatic cysts in sEV-injected mice (Supplemental Figure 7G). Collectively, these data demonstrate that administration of nSMase2-expressing PDA PPT cell sEVs to *KPC; Smpd3^f/f^* mice is sufficient to prevent the prolonged survival phenotype of *KPC; Smpd3^f/f^* mice when compared to *KPC; Smpd3^wt/wt^* mice (Figure 2A) (median survival of *KPC; Smpd3^f/f^* mice injected with sEVs isolated from *KPC; Smpd3^wt/wt^*PDA PPT cell lines is 18.71 weeks vs median survival of *KPC; Smpd3^wt/wt^*mice injected with sEVs isolated from *KPC; Smpd3^wt/wt^* PDA PPT cell lines is 19.43 weeks, Log-rank test p-value = 0.6596), thereby implicating loss of nSMase2-generated exosomes as a mechanism for prolonged survival of *KPC; Smpd3^f/f^* mice.

**Figure 4.**
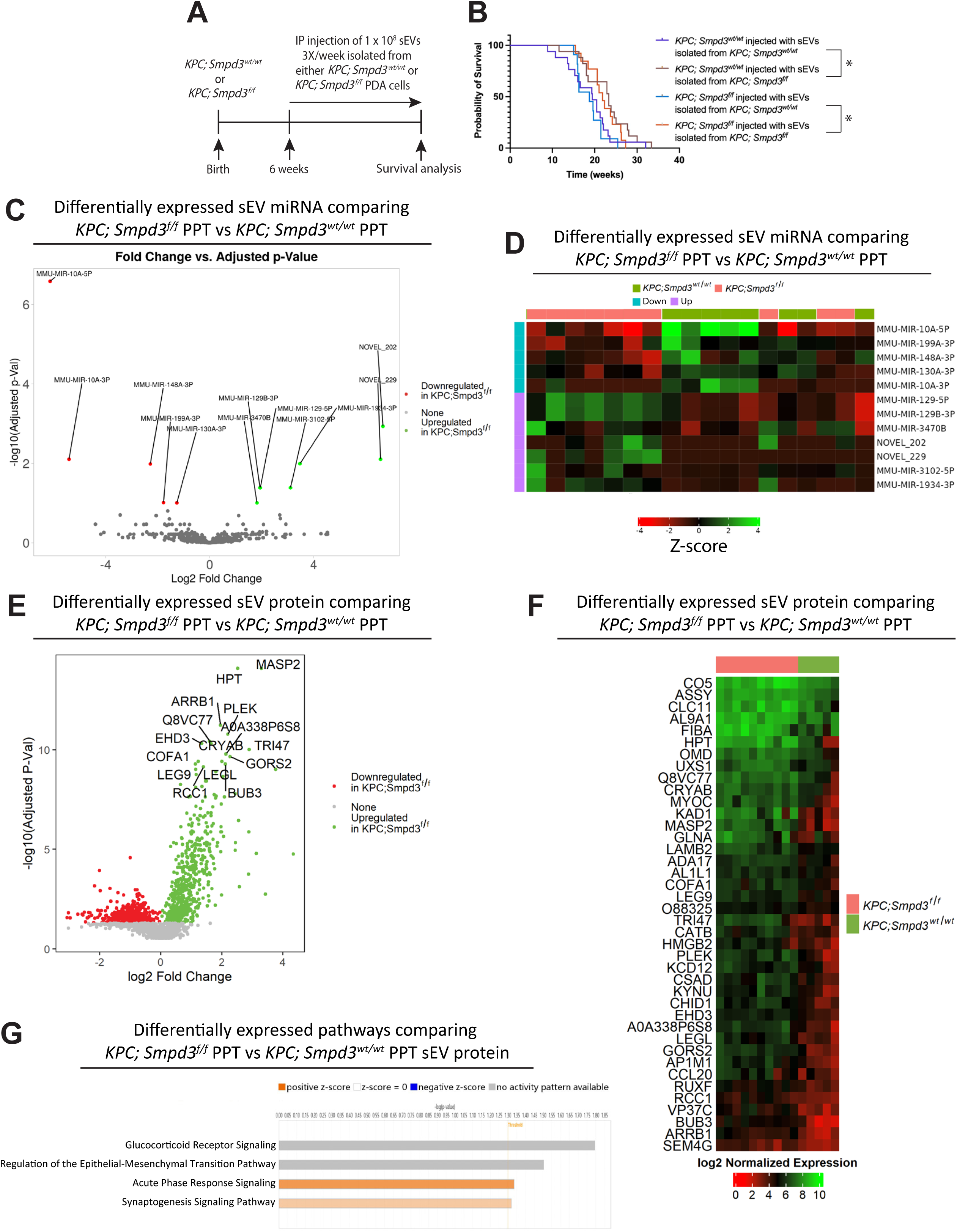
PDA cell exosomes generated through nSMase2 accelerate PDA progression. A) Schematic of exosome injection study. B) Probability of survival for the indicated groups is graphed. Log-rank test was performed between the indicated groups. C-D) Volcano plot (C) and heat map (D) depict differentially expressed exosomal miRNAs isolated from *KPC; Smpd3^f/f^*and *KPC; Smpd3^wt/wt^* polyclonal PDA PPT cell lines. E) Exosomal protein abundance results for *KPC; Smpd3^f/f^*versus *KPC; Smpd3^wt/wt^* PPT cell lines are depicted. The x-axis shows the log2 fold change for *KPC; Smpd3^f/f^* over *KPC; Smpd3^wt/wt^* and the y-axis shows −log10(q-value). Significantly upregulated proteins (q-value < 0.05) are shown in green and significantly downregulated proteins are shown in red. F) Heatmap of top 40 differentially abundant exosomal proteins comparing *KPC; Smpd3^f/f^* versus *KPC; Smpd3^wt/wt^*, displaying log2 normalized expression. Red indicates lower abundance of a given protein in each sample and green indicates higher abundance. Rows and columns are arranged based on hierarchical clustering dendrograms (not displayed). The color bar at the top indicates the genotype. G) IPA results depict significantly differentially regulated pathways when comparing exosomal proteins from *KPC; Smpd3^f/f^* versus *KPC; Smpd3^wt/wt^* PPT cell lines.

### nSMase2 organizes packaging of miRNA and proteins into PDA cell exosomes

The sEV injection study suggests that nSMase2-generated exosome cargo affects PDA progression. Previous studies have demonstrated that exosomes are highly enriched in small RNA species and proteins that can be transferred to and functional in recipient cells (Davis, 2007; Kosaka et al., 2010; Mittelbrunn et al., 2011). To define sEV content, we next performed small RNA-Seq and proteomics analyses on sEVs isolated from the murine *KPC; Smpd3^wt/wt^* and *KPC; Smpd3^f/f^* PDA PPT cell lines (Supplemental Table 4). No significantly differentially expressed PIWI-interacting RNAs (piRNAs) were found. Twelve significantly differentially expressed microRNAs (miRNAs) were identified when comparing sEV miRNA isolated from *KPC; Smpd3^f/f^* PPT cell lines versus *KPC; Smpd3^wt/wt^* PPT cell lines (Figure 4C-D and Supplemental Table 4). MMU-MIR-10A-5P and MMU-MIR-10A-3P are 70.54- and 42.63-fold, respectively, downregulated in sEVs isolated from the murine *KPC; Smpd3^f/f^* PPT cell lines when compared to *KPC; Smpd3^wt/wt^* PPT cell lines. MMU-MIR-10A-5P and MMU-MIR-10A-3P have previously been reported to reduce gene expression signatures associated with proinflammatory macrophage polarization (Cho et al., 2019; Kiran et al., 2023).

One thousand twenty-six significantly differentially expressed (q-value <0.05) proteins were identified when comparing sEV protein isolated from *KPC; Smpd3^f/f^* PPT versus *KPC; Smpd3^wt/wt^* PPT cell lines (Figure 4E-F, Supplemental Table 4). *Smpd3* was significantly downregulated, while *Smpd1* was significantly upregulated in sEVs isolated from *KPC; Smpd3^f/f^*

PPT cell lines when compared to sEVs isolated from *KPC; Smpd3^wt/wt^*PPT cell lines. Components of ESCRT-0 and ESCRT-I complexes including *Stam*, *Mvb12a*, *Vps28*, *Vps37b*, *Vps37c*, and *Tsg101* in addition to the accessory protein ALIX encoded by *Pdcd6ip* were significantly upregulated in sEVs isolated from *KPC; Smpd3^f/f^* PPT cell lines when compared to sEVs isolated from *KPC; Smpd3^wt/wt^* PPT cell lines. In contrast, *Chmp2a*, a component of the ESCRT-III complex, and *Rab35*, a regulator of exosome secretion (Hsu et al., 2010), were significantly downregulated in sEVs isolated from *KPC; Smpd3^f/f^* PPT cell lines when compared to sEVs isolated from *KPC; Smpd3^wt/wt^* PPT cell lines. Glypican-1 was detected in the sEVs but was not significantly different in abundance when comparing sEV protein isolated from *KPC; Smpd3^f/f^*PPT versus *KPC; Smpd3^wt/wt^* PPT cells lines (data not shown).

nSMase2 promotes the packaging of keratin and claudin proteins into exosomes as evident by a significant downregulation of *Krt1*, *Krt2*, *Krt7*, *Krt8*, *Krt18*, *Krt19, Cldn3, Cldn4,* and *Cldn7* in sEVs isolated from *KPC; Smpd3^f/f^* PPT cell lines when compared to sEVs isolated from *KPC; Smpd3^wt/wt^* PPT cell lines. In contrast, the cadherin proteins encoded by *Cdh1*, *Cdh6*, *Cdh11*, *Cdh15*, and *Cdh17* are significantly upregulated in sEVs isolated from *KPC; Smpd3^f/f^* PPT cell lines when compared to sEVs isolated from *KPC; Smpd3^wt/wt^* PPT cell lines. Recruitment of E-cadherin to EVs has previously been shown to depend on the ESCRT-I protein Tsg101 (Banfer et al., 2022). The top two proteins enriched in sEVs isolated from *KPC; Smpd3^f/f^* PPT cell lines, representing sEVs packaged independently of nSMase2, were creatine kinase b (20.27-fold enrichment) and adenylate kinase 1 (13.59-fold enrichment), encoded by *Ckb* and *Ak1*, respectively, enzymes involved in energy metabolism (Supplemental Table 4). The sEV protein differential expression list was used in IPA. Acute Phase Response Signaling and Synaptogenesis Signaling Pathway were significantly upregulated in sEVs isolated from *KPC; Smpd3^f/f^* PPT cell lines compared to sEVs isolated from *KPC; Smpd3^wt/wt^* PPT cell lines (Figure 4G). Together, these data demonstrate that the nSMase2-mediated exosome biogenesis pathway is a critical regulator of PDA cell exosomal microRNA and protein packaging.

### nSMase2 loss results in activation of the immune response in KPC mouse pancreata

Multiple IPA pathways involved in regulating immune responses were differentially expressed in the RNA-seq dataset. When comparing *KPC; Smpd3^f/wt^*PPT cell lines vs *KPC; Smpd3^wt/wt^*PPT cell lines, Dendritic Cell Maturation, IL-33 Signaling Pathway, fMLP Signaling in Neutrophils, Neutrophil Extracellular Trap Signaling Pathway, CD28 Signaling in T Helper Cells, and Natural Killer Cell Signaling were significantly upregulated in *KPC; Smpd3^f/wt^* cell lines (Supplemental Table 2). When comparing *KPC; Smpd3^f/f^* PPT cell lines vs *KPC; Smpd3^wt/wt^* PPT cell lines, IL-13 Signaling Pathway and IL-17 Signaling (McAllister et al., 2014) were significantly downregulated in *KPC; Smpd3^f/f^* PPT cell lines (Supplemental Table 2). IL1, IL1A, and IL6 were predicted to be inhibited in *KPC; Smpd3^f/f^* PPT cell lines when compared to *KPC; Smpd3^wt/wt^*PPT cell lines in IPA Upstream Regulators analysis (Supplemental Table 2).

To investigate whether loss of pancreatic epithelial nSMase2 in KPC mice affects immune cell recruitment in the pancreatic microenvironment, we next quantified immune cells in *KPC; Smpd3^wt/wt^*and *KPC; Smpd3^f/f^* pancreata using a spectral flow cytometry panel. The pancreas from 10-11-week-old *KPC; Smpd3^wt/wt^* and *KPC; Smpd3^f/f^* mice was digested with collagenase and stained using a 17-marker panel. We observed a significant increase in CD45^+^ immune cells (1.570% ± 0.8863 for *KPC; Smpd3^wt/wt^* and 67.63% ± 3.284 for *KPC; Smpd3^f/f^*, p<0.0001), macrophages with a proinflammatory phenotype gated by CD45^+^F4/80^+^Tnfα, cytotoxic T cells gated by CD45^+^CD90.2^+^CD8a^+^, and anti-tumorigenic N1-like neutrophils (Wang et al., 2021) gated by CD45^+^Ly6G^+^Tnfα in *KPC; Smpd3* pancreata when compared to *KPC; Smpd3^wt/wt^* pancreata (Figure 5A-D). A modest increase in CD45^+^F4/80^+^CD11c^+^ macrophages, another gating strategy for the proinflammatory phenotype (Qian et al., 2021), was observed in *KPC; Smpd3^f/f^* pancreata when compared to *KPC; Smpd3^wt/wt^* pancreata (Figure 5D). t-distributed stochastic neighbor embedding (t-SNE) of 200,683 cells stained with the spectral flow panel shows 12 clusters identified for *KPC; Smpd3^wt/wt^* and *KPC; Smpd3^f/f^*pancreata. The heat map of marker expression depicts 2 clusters with scaled median expression of CD45 above 0.5 (Figure 5B-C). These observations suggest that *Smpd3* expression in KPC cells contributes to an immunosuppressive pancreatic microenvironment at preneoplastic stages.

**Figure 5.**
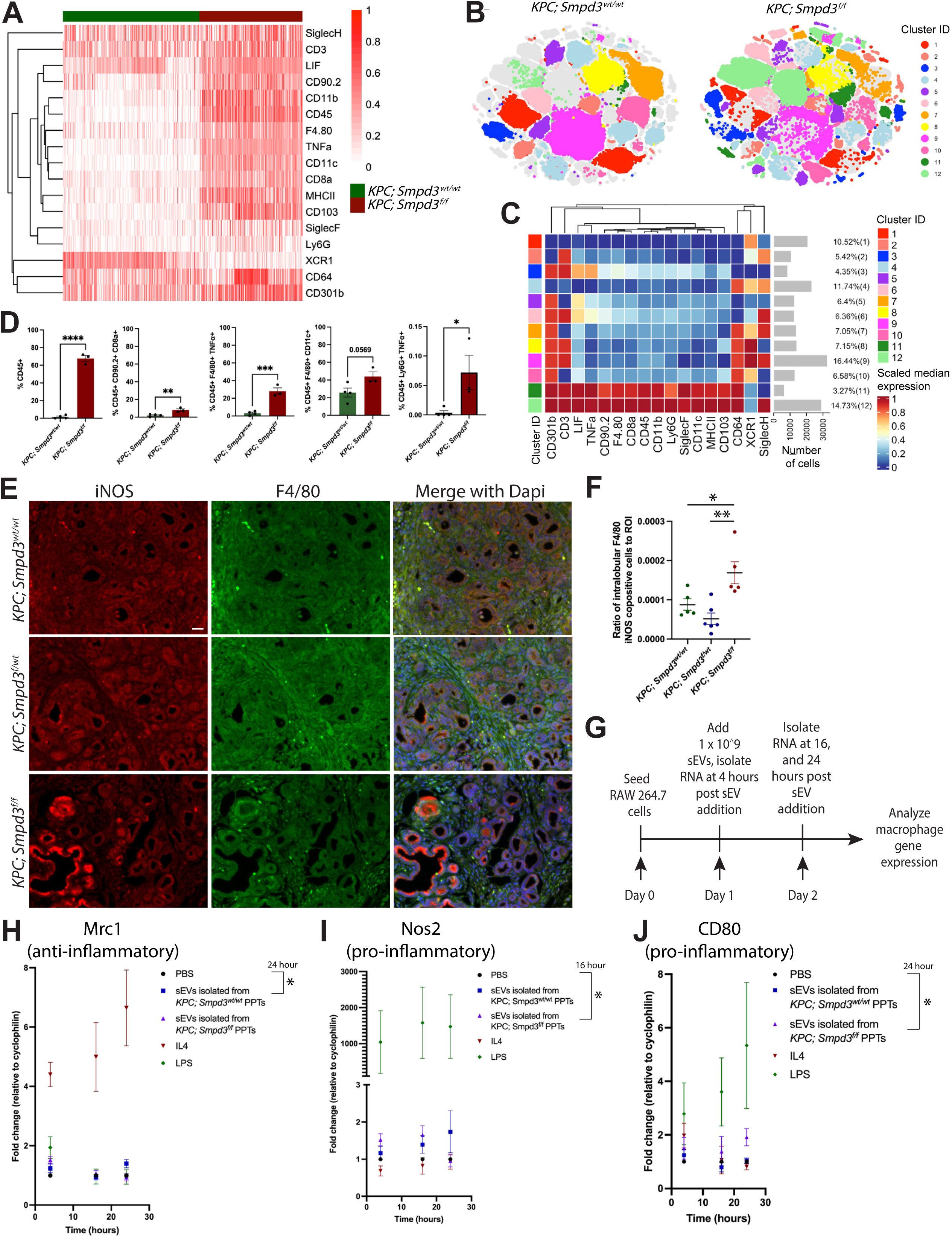
sEVs generated independently of nSMase2 promote a proinflammatory macrophage phenotype. A) Heat map depicts scaled marker expression in *KPC; Smpd3^wt/wt^* and *KPC Smpd3^f/f^* pancreata using our 17-marker myeloid spectral flow cytometry panel. B) t-SNE of 200,683 cells stained with our spectral flow panel depict 12 clusters identified for *KPC; Smpd3^wt/wt^* and *KPC; Smpd3^f/f^* pancreata through FlowSOM clustering. C) Heat map of scaled marker expression and number of cells for clusters shown in B is displayed. D) Bar graphs depict percent cellular subpopulations from *KPC; Smpd3^wt/wt^*and *KPC; Smpd3^f/f^* pancreata for the indicated markers after gating in FlowJo. E) Immunofluorescence images depict co-labeling with F4/80, iNOS, and Dapi at 10-11 weeks of age. Scale bar is 20uM. F) Quantification of intralobular F4/80^+^ and iNOS^+^ copositive cells in *KPC; Smpd3^wt/wt^*(n=5 animals), *KPC; Smpd3^f/wt^* (n=6 animals), and *KPC; Smpd3^f/f^* (n=5 animals) pancreata at 10-11 weeks of age is depicted. G) Schematic of experiment to assess how nSMase2-mediated exosome biogenesis affects polarization of macrophages. H-J) RNA expression fold change in RAW 264.7 cells, with PBS treated normalized to 1, for each gene at the indicated time point and treatment condition is depicted. Results of unpaired t tests between groups shown at the time point written are depicted.

### PDA cell sEVs generated independently of nSMase2 activate anti-tumorigenic macrophage gene expression programs

The immune profiling analyses indicated a prominent infiltration of proinflammatory macrophages in *KPC; Smpd3^f/f^*pancreata. Immunofluorescence staining and quantification for F4/80 demonstrated a significant increase in intralobular, but not interlobular, F4/80^+^ macrophages in *KPC; Smpd3^f/f^* pancreata when compared to *KPC; Smpd3^f/wt^*pancreata at 10-11 weeks of age. Ly6G^+^ intralobular, but not interlobular, neutrophils (Kleinholz et al., 2021), are significantly increased in *KPC; Smpd3^f/f^* pancreata when compared to both *KPC; Smpd3^f/wt^* and *KPC; Smpd3^wt/wt^* pancreata at 10-11 weeks of age (Supplemental Figure 8A-C). Immunofluorescence staining and quantification demonstrated a significant increase in F4/80^+^iNOS^+^ copositive pro-inflammatory macrophages in *KPC; Smpd3^f/f^* pancreata when compared to both *KPC; Smpd3^f/wt^* and *KPC; Smpd3^wt/wt^*pancreata at 10-11 weeks of age (Figure 5E-F). To determine whether nSMase2-generated exosomes affect polarization of macrophages, we next seeded equal numbers of RAW 264.7 cells, then added 1 x 10^9^ sEVs isolated from the polyclonal end-stage *KPC; Smpd3^wt/wt^* or *KPC; Smpd3^f/f^* PPT cell lines, and isolated RNA 4, 16, and 24 hours after sEV addition for assessment of macrophage marker expression (Figure 5G). Macrophages treated with sEVs isolated from *KPC; Smpd3^wt/wt^*PPT cell lines displayed a significant increase in the anti-inflammatory macrophage marker *Mrc1,* that encodes for CD206, when compared to PBS treated macrophages (Figure 5H). We observed a significant increase in the proinflammatory macrophage markers *Nos2*, *CD80*, and *CD86* and a trend towards an increase in the proinflammatory macrophage markers *Il1b* and *Stat1* in macrophages treated with sEVs isolated from *KPC; Smpd3^f/f^* PPT cell lines when compared to PBS treated macrophages (Figure 5I-J and Supplemental Figure 8D-F). Together, these data suggest that PDA cell sEVs generated independently of nSMase2 upregulate expression of proinflammatory macrophage markers in RAW 264.7 cells.

### *SMPD3* expression is an independent prognostic marker for PDA patient outcome

To examine the relationship between the mouse model findings and human disease, we next analyzed *SMPD3* expression in several published PDA patient datasets in addition to PDA patient samples. Analysis of a microarray dataset (Idichi et al., 2017) showed no difference in expression of *SMPD3* when comparing adjacent normal pancreas to primary PDA patient samples (Figure 6A). Analysis of an integrated bulk RNA-seq dataset including samples combined from multiple studies (Chan-Seng-Yue et al., 2020; O’Kane et al., 2020) showed a significant reduction in *SMPD3* expression in metastatic PDA when compared to primary PDA (Figure 6B).

**Figure 6.**
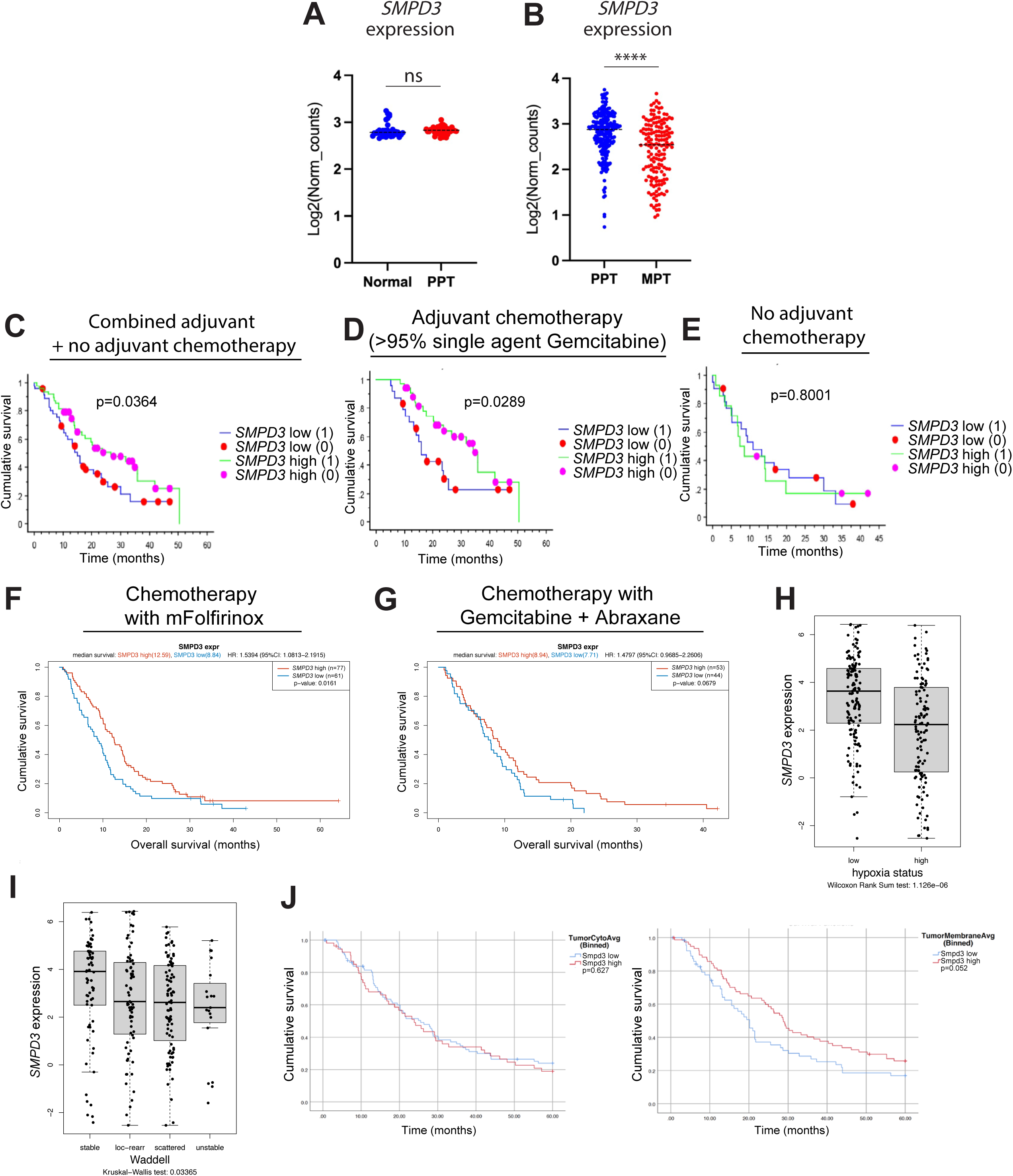
*SMPD3* is an independent prognostic factor for pancreatic cancer patient survival. A) Log2 of the normalized counts are plotted for normal human pancreatic *SMPD3* expression (n=39 samples) and primary PDA *SMPD3* expression (n=39 samples). B) Log2 of the normalized counts are plotted for PPT *SMPD3* expression (n=231 samples) and MPT *SMPD3* expression (n=158 samples) in PDA patients. C) Kaplan-Meier survival curve of the whole cohort (N = 94) with the patients dichotomized into *SMPD3* high and low expression based on the median mRNA expression value. *SMPD3* high group had significantly better survival (median survival of 26.5 vs 15.0 months; P = 0.0364). A (0) value indicates a censored event and a (1) value indicates a death event. D) Kaplan-Meier survival curve of the subset of the whole cohort of patients who underwent adjuvant chemotherapy (> 95% single agent gemcitabine) after surgical resection (N = 58). *SMPD3* expression was significantly associated with better survival (median survival of 34.3 vs 15.9 months; P = 0.0289) indicating *SMPD3* is a biomarker of adjuvant gemcitabine response. E) Kaplan-Meier survival curve of the subset of the whole cohort of patients who did not undergo adjuvant chemotherapy after surgical resection (N = 36). *SMPD3* expression was not associated with survival (P = 0.8001). F-G) Kaplan-Meier curves depict patient survival based on *SMPD3* expression in locally advanced primaries or metastatic PDA from the COMPASS trial for patients who received chemotherapy with modified Folfirinox or Gemcitabine plus Abraxane. *SMPD3* expression was stratified into high and low with low expression defined as the maximal chi-squared statistic. H) Log2 of *SMPD3* expression by hypoxia status defined by Connor et al. 2019. I) Log2 of *SMPD3* expression in defined Waddell structural variation subtypes in COMPASS cases is depicted. J) Kaplan-Meier curves depict patient survival based on cytoplasmic and membranous nSMase2 expression in treatment naïve primary resected PDA. Tumor, and not stromal, cell nSMase2 expression was scored in 143 patients. When comparing survival of PDA patients with low cytoplasmic nSMase2 expression (n=89, mean survival=30.1 months) and high cytoplasmic nSMase2 expression (n=54, mean survival=29.1 months), p=0.627 using the log-rank test. When comparing survival of PDA patients with low membrane nSMase2 expression (n=65, mean survival=25.6 months) and high membrane nSMase2 expression (n=78, mean survival=32.9 months), p=0.052 using the log-rank test.

Analysis of a bulk RNA-seq dataset with high epithelial content (≥40%) (Bailey et al., 2016) demonstrated that high *SMPD3* expression in surgically resected, treatment naïve primary PDA significantly correlated with longer patient survival when patients received adjuvant chemotherapy, more than 95% of which was single agent gemcitabine (Figure 6C-D). No difference in survival was observed in patients who did not receive adjuvant chemotherapy (Figure 6E). Patients who did not receive adjuvant chemotherapy were likely performing too poorly to tolerate or merit it, potentially due to post-operative complications or rapid cancer progression; thus, we would not necessarily expect a difference in survival based on gene expression in this group. Rather, we stratified the dataset based on treatment with adjuvant chemotherapy to remove a variable since nSMase2 expression is known to be affected by multiple chemotherapeutics (Ito et al., 2009; Kong et al., 2015; Shamseddine et al., 2015). IPA analysis of this dataset comparing PPTs with high *SMPD3* expression to PPTs with low *SMPD3* expression that received adjuvant chemotherapy showed pathways regulating metabolism of drugs to be upregulated in PPTs with high *SMPD3* expression including Phase 1 – Functionalization of Compounds and Phase 2 – Conjugation of Compounds (Supplemental Figure 9A and Supplemental Table 5). IPA analysis comparing PPTs with high *SMPD3* expression to PPTs with low *SMPD3* expression that did not receive adjuvant chemotherapy showed pathways regulating ECM organization to be upregulated in PPTs with low *SMPD3* expression (Supplemental Figure 9B and Supplemental Table 5).

Subsequently, we analyzed patient survival based on *SMPD3* expression in locally advanced primaries and metastatic tissue using RNA-seq data from the COMPASS trial, which was enriched for epithelial content by laser capture microdissection (Aung et al., 2018). PDA patients with high *SMPD3* expression who received chemotherapy with mFolfirinox lived significantly longer than PDA patients with low *SMPD3* expression (Figure 6F). We observed a trend (p=0.0679) toward prolonged survival of PDA patients with high *SMPD3* expression who received chemotherapy with Gemcitabine and Abraxane when compared to PDA patients with low *SMPD3* expression (Figure 6G). These data suggest that *SMPD3* expression in both primary PDA and advanced lesions is an independent prognostic marker for PDA patient survival. Using a validated 76 gene hypoxia signature (Connor et al., 2019) (Harris et al., 2015), we found PDA patients in the COMPASS trial with a high hypoxia status have significantly lower *SMPD3* expression than PDA patients in the COMPASS trial with a low hypoxia status (Figure 6H). PDA has previously been categorized into four subtypes of chromosomal structural variation: stable, locally rearranged, scattered, and unstable (Waddell et al., 2015). PDA patients in the COMPASS trial with the stable subtype showed significantly higher *SMPD3* expression when compared to all other subtypes (Figure 6I). These analyses suggest that PDA patients with low *SMPD3* expression have significantly higher hypoxia and more chromosomal aberrations than PDA patients with high *SMPD3* expression, which may be contributing to reduced response to chemotherapeutics and shorter survival.

nSMase2 is localized to multiple compartments of the cell including the plasma membrane and Golgi apparatus (Hostetler and Yazaki, 1979; Tani and Hannun, 2007). To determine if the prognostic significance of nSMase2 expression is dependent on subcellular localization, we scored a PDA patient TMA consisting of resected treatment naïve primary PDA tissue for tumor cell nSMase2 expression and localization. High membranous nSMase2 expression was near-significantly correlated with longer patient survival (p=0.052) when compared to low membranous nSMase2 expression. We observed no correlation between high and low cytoplasmic nSMase2 expression and PDA patient survival (Figure 6J and Supplemental Figure 9C). These data suggest that levels of membranous nSMase2 play an important role in predicting PDA patient outcome.

### nSMase2 mediates PDA vasculature remodeling

When examining the intracellular bulk RNA-sequencing dataset of polyclonal murine PPT cell lines, multiple pathways regulating blood vessel formation, function, and signaling were identified as significantly deregulated including VEGF signaling, VEGF Family Ligand-Receptor Interactions, Apelin Endothelial Signaling Pathway (Helker et al., 2020), Atherosclerosis Signaling, and HIF1α Signaling (Supplemental Table 2 and Figure 7A-B). IPA upstream regulator analysis predicted *Vegfa* to be inhibited in *KPC; Smpd3^f/f^* PPT when comparing *KPC; Smpd3^f/f^* and *KPC Smpd3^f/wt^* (Supplemental Table 2 and Figure 7C). Based on these IPA analysis results and an increased hypoxia status in PDA patients with low *SMPD3* expression, we hypothesized that PDA cell nSMase2 affects vasculature formation; this could explain the opposite Kaplan-Meier curve observations between the mouse models and PDA patient analyses as vasculature abundance and integrity affect chemotherapeutic response, and the mouse models did not receive chemotherapy while the PDA patients did. We next assessed blood vessel abundance by CD31 immunolabeling in the mouse models and human tissue. *KPC; Smpd3^f/f^* mice have significantly fewer CD31^+^ pancreatic endothelial cells than *KPC; Smpd3^wt/wt^* mice at 19-21 weeks of age (Figure 7D-E). Rag1 KO mouse pancreases injected orthotopically with *KPC* LucFlag *Smpd3* shRNA 1 or *KPC* LucFlag *Smpd3* shRNA 2 cell lines have significantly fewer CD31^+^ pancreatic endothelial cells than Rag1 KO mice injected orthotopically with *KPC* LucFlag scrambled shRNA cells 21 days post injection (Figure 7F-G). To assess vessel permeability, we injected *KPC; Smpd3^wt/wt^*and *KPC; Smpd3^f/f^* mice with a high molecular weight dextran at 10-11 weeks of age. *KPC; Smpd3^f/f^* pancreases have significantly higher dextran per field of neoplastic area when compared to *KPC; Smpd3^wt/wt^* pancreases, suggesting increased blood vessel permeability in *KPC; Smpd3^f/f^* pancreases (Supplemental Figure 9D-E). Leaky vasculature has been positively correlated with poor tumor perfusion, increased hypoxia, and ineffective drug delivery in pancreas cancer (Mortezaee et al., 2023). To assess if nSMase2 expression affects chemosensitivity of PDA cells, we determined the IC50 of the polyclonal end-stage murine PDA cell lines. We observed no significant difference in IC50 when comparing *KPC; Smpd3^wt/wt^* vs *KPC; Smpd3^f/f^* PPT cell lines and *KPC* Mock OE vs *KPC Smpd3* OE cell lines (Supplemental Figure 9F).

**Figure 7.**
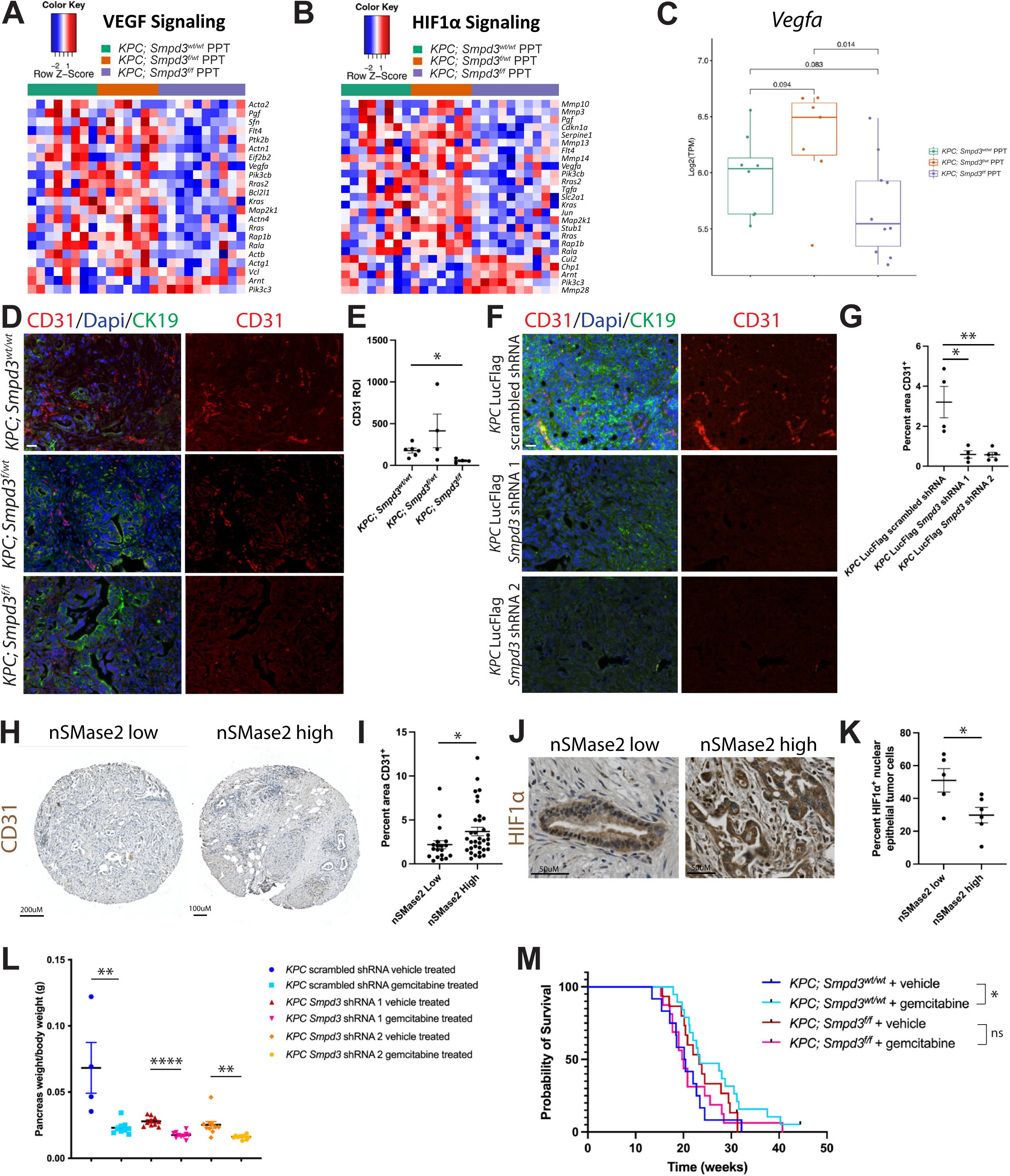
nSMase2 regulates PDA vasculature development. A-B) Heat maps showing VEGF Signaling and HIF1α Signaling pathways in all three PPT cell lines. C) Log of transcript per million (TPM) values for *Vegfa* gene are graphed for sequenced PPT cell lines. Wilcoxon test was performed. D) Pancreatic CD31 and CK19 immunolabeling at 19-21 weeks of age in *KPC; Smpd3^wt/wt^*, *KPC; Smpd3^f/wt^*, and *KPC; Smpd3^f/f^* mice is depicted. Scale bar is 20uM. E) Quantification of pancreatic CD31 immunofluorescence in *KPC; Smpd3^wt/wt^*, *KPC; Smpd3^f/wt^*, and *KPC; Smpd3^f/f^* mice is shown. F) Pancreatic CD31 and CK19 immunolabeling at 21 days post orthotopic injection of *KPC* LucFlag scrambled shRNA, *KPC* LucFlag *Smpd3* shRNA 1, and *KPC* LucFlag *Smpd3* shRNA 2 cell lines into Rag1 KO mice 21 days post injection is depicted. Scale bar is 20uM. G) Quantification of pancreatic CD31 immunofluorescence in Rag1 KO mice injected with *KPC* LucFlag scrambled shRNA, *KPC* LucFlag *Smpd3* shRNA 1, and *KPC* LucFlag *Smpd3* shRNA 2 cell lines 21 days post injection is depicted. H) Immunohistochemistry of CD31 in primary tumors from PDA patients is depicted with low and high nSMase2 expression. I) Percent CD31 positive area in primary nSMase2 low (n=19) and nSMase2 high (n=34) pancreatic tumors from PDA patients is shown. J) Immunohistochemistry of HIF1α is depicted in primary tumors from PDA patients with low and high nSMase2 expression. K) Percent nuclear HIF1α epithelial tumor cells in primary nSMase2 low (n=5) and nSMase2 high (n=6) PDA patient pancreatic tumors is shown. L) Ratio of pancreas weight to body weight for gemcitabine and vehicle treated Rag1 KO mice 21 days after pancreatic orthotopic injection of the depicted cell lines is shown. For *KPC* scrambled shRNA vehicle treated mice, n=4, for *KPC* scrambled shRNA gemcitabine treated mice, n=9, for *KPC Smpd3* shRNA 1 vehicle treated mice, n=10, for *KPC Smpd3* shRNA 1 gemcitabine treated mice, n=10, for *KPC Smpd3* shRNA 2 vehicle treated mice, n=10, and for *KPC Smpd3* shRNA 2 gemcitabine treated mice, n=10. M) Kaplan-Meier survival curves show probability of survival for vehicle-injected *KPC; Smpd3^wt/wt^*mice (n=12, median survival=20.29 weeks), gemcitabine-injected *KPC; Smpd3^wt/wt^* mice (n=19, median survival=23.43 weeks), vehicle-injected *KPC; Smpd3^f/f^* mice (n=15, median survival=23.29 weeks), gemcitabine-injected *KPC; Smpd3^f/f^* mice (n=16, median survival=19.86 weeks). A black line indicates a censored animal.

PDA patients with low nSMase2 expression in the PPT have significantly fewer CD31^+^ pancreatic endothelial cells than PDA patients with high nSMase2 expression in the PPT (Figure 7H-I). PDA patients with low nSMase2 expression also express significantly higher levels of nuclear, but not cytoplasmic, HIF1α when compared to PDA patients with high nSMase2 expression in the PPT (Figure 7J-K, Supplemental Figure 9G). To determine if nSMase2 expression affects chemotherapeutic response *in vivo*, we next treated the mouse models with gemcitabine. Rag1 KO mouse pancreases injected orthotopically with *KPC* LucFlag *Smpd3* shRNA 1 and *KPC* LucFlag *Smpd3* shRNA 2 cell lines show an average of 1.59- and 1.57-fold reduction in pancreas weight/body weight ratio, respectively, for mice treated with gemcitabine when compared to vehicle treated controls. Rag1 KO mouse pancreases injected orthotopically with the *KPC* LucFlag scrambled shRNA cell line show an average of 2.97-fold reduction in pancreas weight/body weight ratio for mice treated with gemcitabine when compared to vehicle treated controls (Figure 7L). Additionally, *KPC; Smpd3^wt/wt^* mice respond to chemotherapy with single agent gemcitabine (median survival of vehicle-injected *KPC; Smpd3^wt/wt^* mice is 20.29 weeks vs median survival of gemcitabine-injected *KPC; Smpd3^wt/wt^* mice is 23.43 weeks, p=0.0272), while *KPC; Smpd3^f/f^* mice do not (median survival of vehicle-injected *KPC; Smpd3^f/f^* mice is 22.64 weeks vs median survival of gemcitabine-injected *KPC; Smpd3^f/f^* mice is 19.86 weeks, p=0.4972) (Figure 7M). These data indicate that pancreatic tumors with reduced *Smpd3* expression have a reduced response to chemotherapy when compared to controls. Importantly, when comparing Kaplan-Meier data of gemcitabine-injected *KPC; Smpd3^wt/wt^* mice (median survival 23.43 weeks) to gemcitabine-injected *KPC; Smpd3^f/f^* mice (median survival 19.86 weeks), the p-value is 0.0477 using the log-rank test; these data demonstrate that treatment of the *KPC; Smpd3^wt/wt^* and *KPC; Smpd3^f/f^* mouse models with gemcitabine reverses the Kaplan-Meier curve results shown in Figure 2A to match the correlation seen between *SMPD3* expression and survival in Kaplan-Meier analyses of PDA patients who also received chemotherapy (Figure 6). Together, these data suggest that reducing pancreatic nSMase2 expression hinders vasculature development, promotes formation of hypoxia, and reduces chemotherapeutic response.

## DISCUSSION

The concept of tumor cell extrinsic factors, particularly exosomes, mediating cell-cell communication within the TME to alter tumorigenesis has been of great interest to researchers. Our study manifested a role for PDA cell exosomes generated through the ceramide-dependent pathway in the context of mutant Kras and heterozygous p53 loss in driving pancreatic tumor growth. Epithelial nSMase2 loss in KPC mouse pancreata delays preneoplasia and neoplasia development, promotes the formation of a molecularly distinct tumor subtype, and abrogates vasculature development, which likely reduces chemotherapeutic response in PDA patients and KPC mice with diminished nSMase2 expression.

A multitude of transcriptomic and lipidomic changes were unveiled by altering *Smpd3* expression in KPC cells, suggesting nSMase2-generated ceramide and/or phosphocholine have substantial intrinsic effects in modifying PDA cell biology *in vitro*. The orthotopic injection study of PDA cells with reduced *Smpd3* expression into Rag1 KO mice resulted in slowed tumor growth when compared to controls, suggesting that potential mature B and T cell infiltration and function are not required for the delayed tumor growth phenotype. While reduced *in vivo* proliferative capacity could partly explain reduced malignant potential with *Smpd3* knockdown in PDA cells, we observed no difference in proliferation *in vitro* when comparing *Smpd3* knockdown murine PDA cells with the control. These data rule out a prominent role for autocrine effects of PDA cell exosomes generated through nSMase2 in regulating PDA cell proliferation *in vitro* and suggest that the TME is a major player in mediating the delayed growth phenotype of *Smpd3* knockdown PDA cells in Rag1 KO mice. Contrarily, previous work has demonstrated abrogating exosome biogenesis machinery by knocking down Rab27A and Rab27B reduced the proliferative and invasive capacity of human PDA cells, highlighting diverse roles for exosome biogenesis pathways in regulating PDA cell growth (Li et al., 2017).

Our findings could potentially be explained by interactions between nSMase2-generated exosomes and TME milieu. Epithelial nSMase2 loss in KPC mouse pancreata results in a strong immune response that includes pancreatic infiltration of cytotoxic T cells, N1-like anti-tumorigenic neutrophils, and macrophages with a proinflammatory phenotype during preneoplasia development. Recruited proinflammatory macrophages play a crucial role in initiating formation of pancreatic lesions (Liou et al., 2015; Liou et al., 2013) and have well-known anti-cancer effects in established tumors. Recruited anti-inflammatory, tumor-promoting macrophages promote an immunosuppressive microenvironment and fibrogenesis during early and late stage PDA (Bastea et al., 2019; Liou et al., 2017). Our observations that sEVs generated from nSMase2-expressing PDA cells upregulate the anti-inflammatory macrophage marker *Mrc1* in RAW 264.7 cells, and sEVs generated independently of nSMase2 upregulate expression of several proinflammatory macrophage markers in RAW 264.7 cells demonstrate a functional role for the ceramide-dependent exosome biogenesis pathway in macrophage polarization. A possible mechanism is that MMU-MIR-10A-5P and MMU-MIR-10A-3P are upregulated in sEVs generated through nSMase2, which have previously been reported to reduce gene expression signatures associated with proinflammatory macrophage polarization (Cho et al., 2019; Kiran et al., 2023). Pancreatic KPC cell exosomes generated independently of nSMase2 might potentiate macrophage polarization toward proinflammatory in *KPC; Smpd3^f/f^* pancreata. The enhanced immune infiltration in *KPC; Smpd3^f/f^* mice at 10-11 weeks of age (pre-tumor) may contribute to an overall anti-tumorigenic microenvironment that delays neoplastic growth.

The delayed neoplasia development and reduced tumor burden in *KPC; Smpd3^f/f^* mice is accompanied by defective vasculature development, leading to a model that responds poorly to chemotherapy; this observation is mirrored by the patient dataset analyses with reduced nSMase2 expression in PDA tissues correlating with decreased CD31^+^ blood vessels, hypoxia and decreased response to chemotherapy as measured by survival. Intratumoral hypoxia and compromised blood vessel integrity are widely established to contribute to PDA treatment barriers and poor survival (Ansardamavandi et al., 2023; Liu et al., 2019; Tao et al., 2021). Our observation is in line with previous reports showing that nSMase2-generated cancer cell exosomes stimulate angiogenesis (Kosaka et al., 2013; Shang et al., 2020).

The mouse model data collectively implicate nSMase2-expressing PDA cell sEVs in promoting PDA development and PDA cell sEVs generated independently of nSMase2 in delaying PDA progression. Support for this notion is derived from the exosome injection study using KPC mice. Our study warrants further investigation into the potential therapeutic relevance of targeting specific exosome biogenesis pathways in combination with chemotherapy to combat pancreatic cancer.

## MATERIALS AND METHODS

### Lentiviral constructs

*Smpd3* shRNA1 (TRCN0000099416) and *Smpd3* shRNA2 (TRCN0000099418) were purchased from Sigma-Aldrich and cloned into MISSION^®^ pLKO.1-puro (Sigma-Aldrich, SHC001). The control pLKO.1-scrambled shRNA vector was purchased from Addgene (Plasmid 136035). Mouse *Smpd3* cDNA (GenBank accession number, NM_021491.4) was amplified from a cDNA library generated from a *KPC; Smpd3^wt/wt^* primary murine pancreatic cancer cell line and cloned into pLenti-CMV-EGFP-Blasticidin lentiviral vector (Addgene, 17445) after removal of eGFP region and switching blasticidin for puromycin.

Firefly luciferase cDNA was amplified from pGL3 (Promega) and subcloned into pLenti-CMV-EGFP-Blasticidin lentiviral vector (Addgene, 17445) after removal of eGFP region. Mouse CD63 cDNA (GenBank accession number, NM_001042580) was amplified from a cDNA library generated from the *KPC; Smpd3^wt/wt^* 14 primary murine pancreatic cancer cell line. Mouse CD63 (without stop codon) and eGFP cDNA (without 1^st^ ATG) were linked using 4 amino acid “DLEL” and subcloned into pLenti-CMV-EGFP-Blasticidin lentiviral vector (Addgene, 17445) backbone. pCT-CD63-GFP was purchased from System Biosciences (CYTO120-PA-1). Cell lines used for subcutaneous injections expressed mCherry. mCherry was amplified from MP-283: pSicoR-BstXI-EF1a-puro-T2A-mCherry, cloned into pLenti-CMV-EGFP-Blasticidin lentiviral vector (Addgene, 17445) in the place of eGFP region, and introduced into the cell line via viral transduction and selection. All constructs were verified by sequencing.

Stable cell lines expressing shRNAs and/or cDNAs were established by lentiviral mediated gene transfer. Briefly, cells were infected with viral particles generated by UCSF ViraCore. Infected cells were selected for with puromycin or blasticidin. For generation of cell lines expressing more than one lentiviral construct, lentiviral vectors expressing CD63, flag-tagged firefly luciferase, or mCherry were transduced and selected first. Then, plasmids expressing shRNAs were transduced and selected.

### Immunocytochemistry

Organoids were grown in chamber slides (CELLTREAT Scientific Products, 229168). Cells were washed with 1X PBS-/- (Gibco, 14190-144) (prewarmed at 37°C) for at least 20 minutes, fixed in 4% PFA (prewarmed at 37°C) for 30 minutes at 37°C, then washed three times for 10 minutes each in 1X PBS-/- at room temperature. Organoids were blocked in 10% FBS (Corning, 35011CV), 0.5% Triton X-100 (Sigma-Aldrich, T8787) in 1X PBS-/- overnight at 4°C. Block was aspirated. Primary antibody was diluted in blocking buffer, added to cells, and incubated at 4°C for 48 hours. Cells were washed 3 times for 10 minutes each in 1X PBS-/-. Secondary antibody was diluted in blocking buffer and incubated overnight at 4°C. Cells were washed three times for 10 minutes each in 1X PBS-/-. Cells were stained with Dapi (Sigma-Aldrich, D9564) at 2.5ug/mL for 15 minutes. Cells were washed three times for 10 minutes each in 1X PBS-/-. Organoids were mounted using ProLong Glass Antifade (Invitrogen, P36984). Slides were stored at 4°C protected from light.

### Histology and immunostaining

Tissues were fixed, processed, embedded, and stained using immunofluorescence and immunohistochemistry as previously described in detail (Hendley et al., 2021). Primary antibodies used in this study are listed in Supplemental Table 6. Secondary antibodies used in this study were obtained from Invitrogen, Jackson ImmunoResearch, and Biotium.

Hematoxylin and eosin (H&E), alcian blue, and picrosirius red staining were performed using standard protocols. For H&E staining, Mayer’s Hematoxylin Solution (Sigma-Aldrich, MHS32) and Eosin Y Solution (Sigma-Aldrich, HT110132) were used. For alcian blue staining, Alcian blue solution (Sigma-Aldrich, B8438) and Nuclear Fast Red (Vector Laboratories, H-3403) were used. For picrosirius red staining, Picro Sirius Red Stain Kit (Abcam, ab150681) and Weigert’s Iron Hematoxylin Set (Sigma-Aldrich, HT1079) were used.

For quantification of epithelial (CK19^+^, Vimentin^-^), proliferative (Ki67^+^) pancreatic cancer cells (Dapi^+^), 4-5 images containing 1,862 - 3,168 total CK19^+^, Vimentin^-^, Ki67^+^, Dapi^+^ cells per animal were analyzed in Fiji. For quantification of epithelial (E-cadherin^+^, Vimentin^-^), apoptotic (cleaved caspase 3^+^) pancreatic cancer cells (Dapi^+^), 4-5 images containing 1,103 – 2,453 total E-cadherin^+^, Vimentin^-^, cleaved caspase 3^+^, Dapi^+^ cells per animal were analyzed in Fiji. Cells were defined as having a Dapi^+^ nucleus in the imaging plane. For nSMase2 expression scoring, 2-4 sections per animal were evaluated. For nSMase2 subcellular localization scoring, criteria used to call absent, cytoplasmic, and membranous was that the staining pattern had to represent >50% of all staining patterns observed for that animal. Alcian blue stained slides were scanned and quantified in QuPath. Whole slide scanning was performed using a Zeiss Axio Scan.Z1 whole slide scanner; images were collected with a Zeiss Plan-Apochromat 20x/0.8NA (WD=0.55mm) M27 objective lens in the brightfield mode with a Hitachi HV-F202 camera. Edema scoring criteria of 1-3 were evaluated by a board-certified pathologist. Quantification of tumor area was performed using scanned H&E images in QuPath. Histopathology of *KPC; Smpd3^wt/wt^*, *KPC; Smpd3^f/wt^*, and *KPC; Smpd3^f/f^* mice was reviewed and classified by a board-certified pathologist. Criteria for classification of undifferentiated carcinomas (malignant neoplasm not showing definitive direction of differentiation with more diffuse sheet-like growth pattern) included sarcomatoid undifferentiated carcinoma (malignant undifferentiated neoplasm displaying at least 80% spindle-shaped neoplastic cells (sarcomatoid cells)) and carcinosarcoma (biphasic malignant neoplasm with distinct epithelioid and spindle-shaped cells, each constituting at least 30% of the tumor). Picrosirius red stained slides were scanned using both brightfield and polarized light microscopy on a Nikon Eclipse TE2000-U and quantified using Fiji. For FAP, αSMA, Vimentin, and Dapi quantification, 2-3 immunofluorescent images per animal were analyzed in Fiji. For Gfap, αSMA, and Desmin quantification, 2 images per animal were analyzed in Fiji. For murine CD31 quantification, 3-6 images per animal were analyzed in Fiji. For F4/80 and Ly6G quantification, 2-4 images per animal were analyzed in Fiji. For F4/80 and iNOS quantification, 3-5 images per animal were analyzed in Fiji. Vascular permeability was assessed by percentage of total neoplastic area that was FITC-dextran positive in Fiji using fresh frozen sections and 3-4 images per animal.

### PDA patient TMA IHC and scoring

The clinically annotated pancreatic cancer tissue microarray (TMA) has been previously described (Manuyakorn et al., 2010). After initial deparaffinization and heat-induced epitope retrieval (10mM Tris-EDTA, pH 8.0) with vegetable steamer, TMA sections were blocked in 5% normal goat serum, incubated overnight at 4°C with 1:100 anti-nSMase2 (sc-166637) and further processed with SignalStain® Boost IHC Detection Reagent (Cell Signaling Technology, #8125) and DAB Substrate Kit (CST #8059) per manufacturer’s instructions. Two surgical pathologists (DWD and IRR) evaluated cytoplasmic, membranous, and staining intensity in each of 3 TMA cores per tumor using a semi-quantitative scoring system (range 0-3, 0-absent staining in >90% of tumor; 1-weak staining in >10% tumor; 2-moderate staining in >50% of tumor; 3-strong staining in greater than 50% of tumor). Only those tumors with tumor identified in cores were included in the final analysis (n=143). For survival analysis, each tumor was dichotomized to either a low or high nSMase2 group for cytoplasmic (average score of ≤1 versus >1) and membrane (average score of 0 versus >0) expression. Overall survival time was calculated as the date of surgery to the date of death due to any cause or last clinical follow up to 60 months. Survival estimates were generated using the Kaplan-Meier method and compared using log-rank tests.

For CD31 and HIF1α IHC, the same TMA scored for cytoplasmic and membrane nSMase2 expression was used. IHC was performed using ImmPRESS Polymer reagents (Vector Laboratories). Percent positive area was calculated in QuPath using a classifier for CD31 and total HIF1α quantification. HIF1α nuclear localization was quantified manually in QuPath counting between 93-245 tumor epithelial cells per patient. The average of cytoplasmic and membrane nSMase2 staining scores per patient was classified as low (<0.5) or high (≥0.5).

### RNA isolation, cDNA library preparation, and qPCR

For qPCR and cellular RNA-sequencing, RNA was isolated from cell lines using the RNeasy Mini Kit (Qiagen, 74106) or the RNeasy Plus Micro Kit (Qiagen, 74034) as per the manufacturer’s instructions. For cellular RNA-seq, cDNA libraries were prepared and sequenced by Novogene. Briefly, messenger RNA was purified from total RNA using poly-T oligo-attached magnetic beads. After fragmentation, the first strand cDNA was synthesized using random hexamer primers. Then the second strand cDNA was synthesized using dUTP, instead of dTTP. The directional library was ready after end repair, A-tailing, adapter ligation, size selection, USER enzyme digestion, amplification, and purification. The library was checked with Qubit and real-time PCR for quantification and bioanalyzer for size distribution detection. Quantified libraries were pooled and sequenced using the Illumina NovaSeq S4 6000 system, according to effective library concentration and data amount. A paired end 150bp sequencing strategy was used; 20M reads were sequenced from each end for a total of >40M raw reads/sample.

For qPCR, cDNA was prepared using the SuperScript III First Strand synthesis kit (Thermo Fisher Scientific, 18080085). qPCR was performed using FastStart Universal SYBR Green mix (Sigma-Aldrich, 4913914001). RNA expression of target genes was normalized to *GAPDH* for human samples and *Ppia* for murine samples. qPCR primer sequences are included in Supplemental Table 6. In this study, qPCR was performed on at least 3 passages of the same cell line (Supplemental Figure 1A-B) or at least 3 independent experiments (Figure 5H-J and Supplemental Figure 8D-F), and data are presented as the average of passages or independent experiments plus minus the standard error of the mean. All qPCR reactions were done using three technical replicates for each gene. qPCR data was analyzed using the 2^−ΔΔCt^ method.

For small RNA-Seq performed on isolated sEVs/exosomes, RNA was isolated using the miRNeasy kit (Qiagen, 217084). cDNA libraries were prepared and sequenced by Novogene. Sample quality was assessed by High Sensitivity RNA Tapestation (Agilent Technologies, 5067-5579 and 5067-5580) and quantified by Qubit RNA HS assay (Thermo Fisher Scientific, Q32855). Library preparation was performed with QIAseq miRNA Library Preparation Kit (Qiagen, 331505) following manufacturer’s instructions. Final library size is 180 bp. Illumina 8-nt single-indices were used for multiplexing. Samples were pooled and sequenced on Illumina NovaSeq 6000 sequencer for 150 bp read length in paired-end mode, with an output of 20 million paired end reads per sample (10M in each direction).

### Analysis of published microarray, RNA-seq, and genomic DNA sequencing datasets

#### ICGC RNA-Seq dataset

The patient and tumor samples (N = 94) included in the analysis in Figure 6C-E are a subset of the International Cancer Genome Consortium (ICGC) PACA-AU cohort, contribution from the Australian Pancreatic Cancer Genome Initiative (APGI), of which underwent full transcriptome and whole genome sequencing. The full cohort description is previously described in detail (Bailey et al., 2016). In short, tumor tissue was acquired from surgically resected Stage 1 and 2 PDA specimens after informed consent was obtained from all subjects following Human Research Ethics Committee approval. Tissue dissection of primary material, RNA and DNA extraction was performed, then whole genome and whole transcriptome sequencing was performed using previously published methods (Bailey et al., 2016). Clinicopathological variables and clinical outcomes were collected as part of the previous study.

#### Analysis of existing RNA-Seq and genomic DNA sequencing datasets

The following publicly available datasets were reanalyzed. Normalized counts data was transformed using EdgeR: log2(CPM+c). Differentially expressed genes between experimental groups were then determined using DESeq2 (version 4.0.3) (Love et al., 2014). Log2 normalized counts expression between normal vs primary (NCBI GSE15471) (Badea et al., 2008), primary vs metastasis (EGAD00001005799 and EGAD00001006081) (Chan-Seng-Yue et al., 2020) (O’Kane et al., 2020) and normal vs primary vs metastasis in human organoids (NCBI dbGaP archive accession number phs001611.v1.p1) (Tiriac et al., 2018) was plotted for the representative genes.

RNA sequencing data of locally advanced primary and metastatic tissue from PDA patients enrolled in the COMPASS trial (EGAD00001009409) (Perera et al., 2023) was used to compare *SMPD3* expression with PDA patient survival and hypoxia status. Briefly, biospecimens underwent laser capture microdissection (LCM) for tumor cell enrichment, and RNA-Seq analysis was performed at the Ontario Institute of Cancer Research as per published standard protocol (O’Kane et al., 2020). Reads were aligned to the human reference genome (hg38) and transcriptome (Ensembl v100) using STAR v.2.7.4a. Picard v. 2.21.4 (https://github.com/broadinstitute/picard) was used for marking duplicated reads. Gene expression was calculated in transcripts per million reads mapped (TPM) using stringtie package v. 2.0.6. Log2 transformed expression was used downstream. For survival analyses, *SMPD3* low and high are defined as the maximal chi-squared statistic. Analysis of hypoxia status was conducted as previously described (Connor et al 2019). Whole genome sequencing data from EGA00001011129, which was processed as described (Golan et al., 2021), from locally advanced primary and metastatic tissue from PDA patients enrolled in the COMPASS trial was analyzed to determine the Waddell structural variation subtypes as previously described (Waddell et al., 2015).

### Analysis of RNA-Seq and small RNA-Seq datasets

#### RNA-Seq processing for IPA and DEG analysis

FASTQ files of raw reads were subjected to Salmon v0.9.1 (Patro et al., 2017) using arguments “--gcBias --seqBias”. Genome Reference Consortium Mouse Build 38 (GRCm38) and RefSeq transcript annotation (GCF_000001635.26) were used to quantify the total expected read counts per gene, as well as normalized gene expression in Transcripts per Million (TPM).

#### IPA and DEG analysis

DESeq2 (Love et al., 2014) was used to identify DE genes using gene counts (adjusted p-value ≤ 0.05). DEGs were subjected to IPA. For analysis of RNA-Seq of mouse PDA cell lines, the following comparisons were performed: *KPC; Smpd3^f/f^* PPT cell lines (n=10 animals) vs *KPC; Smpd3^wt/wt^*PPT cell lines (n=8 animals), *KPC; Smpd3^f/wt^* PPT cell lines (n=7 animals) vs *KPC; Smpd3^wt/wt^* PPT cell lines (n=8 animals), *KPC; Smpd3^f/f^* PPT cell lines (n=10 animals) vs *KPC; Smpd3^f/wt^*PPT cell lines (n=7 animals), *KPC; Smpd3^wt/wt^* MPT cell lines (n=6 animals) vs *KPC; Smpd3^wt/wt^* PPT cell lines (n=8 animals), *KPC; Smpd3^f/f^* MPT cell lines (n=4 animals) vs *KPC; Smpd3^f/f^* PPT cell lines (n=10 animals), *KPC; Smpd3^f/f^*MPT cell lines (n=4 animals) vs *KPC; Smpd3^wt/wt^* MPT cell lines (n=6 animals), and KPC *Smpd3* OE (n=4 animals) vs KPC Mock OE (n=4 animals) (Supplemental Table 2).

For analysis of the patient published dataset (Bailey et al., 2016), the following comparisons were performed: PDA patients with high *SMPD3* in PPTs with adjuvant chemotherapy vs high *SMPD3* in PPTs with no adjuvant chemotherapy, PDA patients with high *SMPD3* in PPTs vs PDA patients with low *SMPD3* in PPTs receiving adjuvant chemotherapy, PDA patients with high *SMPD3* in PPTs vs PDA patients with low *SMPD3* in PPTs receiving no adjuvant chemotherapy, and PDA patients with low *SMPD3* in PPTs with adjuvant chemotherapy vs low *SMPD3* in PPTs with no adjuvant chemotherapy (Supplemental Table 5).

#### Isoform analysis comparing all three PPT groups

The raw fastq files for all samples were processed with kallisto (Bray et al., 2016) version 0.45 using the Ensembl GRCm38 as the reference transcriptome to estimate transcript abundances (bootstrapped quantification, n=100), and the output was processed for differential expression analysis using the sleuth package in R (version 0.30) (Pimentel et al., 2017). We used the likelihood ratio test (LRT) to compare all three PPT groups and identify differentially expressed genes (Supplemental Table 2) by aggregating their corresponding isoform level p-values (using aggregation_column = ‘ens_gene’). Isoform-level estimated counts across all samples were visualized for selected DE genes (*Ccn2*, *Fn1*, *Ly6e*, and *Hmga1*) using the plot_bootstrap function within sleuth, with the most significant protein-coding isoform shown in Supplemental Figure 4B.

#### Consensus clustering (CC) for subtype analysis

Samples were subtyped by the Collisson (3), Bailey (4), and Moffitt (5) schemas. In brief, 62 genes identified by Collisson et al., 613 differentially expressed genes from the multiclass SAM analysis in Bailey et al.’s study, and 50 tumor specific genes from Moffitt et al.’s study were utilized for subtyping analysis, seeking the presence of 3, 4 and 2 clusters respectively. Mouse genes were converted to all uppercase to overlap with earlier mentioned human genes. Unsupervised CC was applied for each of the subtyping schemas using the ConsensusClusterPlus (version 1.56.0) package in R (version 4.1.3) for genes (rows) and samples (columns) separately. For clustering of samples, quantile normalization was performed transcriptome-wise after the gene expression was log2 transformed. A distance matrix was derived based on Pearson correlation. Then CC was applied to this distance matrix, which consisted of 1,000 iterations of k-means clustering using Euclidean distance, with 80% items hold-out at each iteration. For clustering of genes, row-scaling was not applied. A distance matrix was derived based on Pearson correlation. Then CC was applied to this distance matrix, which consisted of 200 iterations of k-means clustering using Euclidean distance, with 80% items hold-out at each iteration. The number of K for genes were determined as the same with the original schemas, namely 3, 4 and 2 for Collisson, Bailey, and Moffitt schemas respectively.

#### Analysis of small RNA-Seq

For small RNA-Seq, raw reads of fastq format were processed using custom perl and python scripts. In this step, clean data (clean reads) were obtained by removing reads containing ploy-N, with 5’ adapter contaminants, without 3’ adapter or the insert tag, containing ploy A or T or G or C and low-quality reads from raw data. At the same time, Q20, Q30, and GC-content of the raw data were calculated. Then, a certain range of length from clean reads was chosen to do all the downstream analyses. The small RNA tags were mapped to reference sequence using Bowtie (Langmead et al., 2009) without mismatch to analyze their expression and distribution on the reference. Mapped small RNA tags were used to identify known miRNA. miRBase20.0 was used as reference. Modified software mirdeep2 (Friedlander et al., 2012) and srna-tools-cli were used to obtain the potential miRNA and draw the secondary structures. Custom scripts were used to obtain the miRNA counts as well as base bias on the first position of identified miRNA with a certain length and on each position of all identified miRNA respectively. To remove tags originating from protein-coding genes, repeat sequences, rRNA, tRNA, snRNA, and snoRNA, small RNA tags were mapped to RepeatMasker, Rfam database or those types of data from the specified species itself. The characteristics of hairpin structure of miRNA precursor was used to predict novel miRNA. The software miREvo (Wen et al., 2012) and mirdeep2 were integrated to predict novel miRNA through exploring the secondary structure, the Dicer cleavage site, and the minimum free energy of the small RNA tags unannotated in the former steps. At the same time, custom scripts were used to obtain the identified miRNA counts as well as base bias on the first position with certain length and on each position of all identified miRNA, respectively. Predicting the target gene of miRNA was performed by miRanda (Enright et al., 2003) for animals. miRNA expression levels were estimated by TPM (transcript per million) using previously described criteria (Zhou et al., 2010).

The small RNA sequences annotated as repeats and the unknown sequences were then used for piRNA analysis. To identify known piRNA, the aforementioned small RNA sequences are first aligned to the piRNABank database using Bowtie; then a k-mer frequency method is applied to predict novel piRNA. The count values of both known and novel piRNA are calculated from uniq-count.

For both miRNA and piRNA analysis, data normalization was carried out using the R package DESeq2 (Love et al., 2014). Unless specified, default parameters were used in the following post-sequencing analysis. Briefly, counts data was transformed using EdgeR: log2(CPM+c). Differentially expressed genes between experimental groups were then determined using DESeq2 (version 4.0.3). An FDR cutoff of 0.1 was used for analysis of small RNA-seq data.

### Analysis of flow cytometry panel

#### Spectral flow data analysis

Spectral flow data were unmixed and compensated using SpectroFlow (Cytek Biosciences, v3.2.1) and further analyzed in FlowJo (verion 10.10.0). For dimensionality reduction, manual gating in FlowJo was performed on cells > singlets > live cells (Zombie NIR^-^). Arcsinh transformation, quality control, and dimensionality reduction of multicolor spectral flow data was then performed in R and essentially as described (den Braanker et al., 2021). Briefly, flow cytometry data was imported to R using the flowCore package (Hahne et al., 2009) and downsampled to achieve equal cell numbers for all samples (28669 cells/sample). Flow data was normalized using the arcsinh cofactor transformation method of the flowVS package (Azad et al., 2016) and automated quality control of transformed data was performed using the PeacoQC package (Emmaneel et al., 2022) before it was converted to a “SingleCellExperiment” object in R. Subsequently, dimensionality reduction, clustering, and visualization of spectral flow data was performed using Seurat, CATALYST (Crowell et al., 2024), dittoSeq, and scater toolkits (Bunis et al., 2021; Hao et al., 2021; McCarthy et al., 2017). T-distributed stochastic neighbor embeddings (t-SNE) and clustering was generated based on all markers included in the spectral flow panel. Raw data and computational pipelines used in this study are available upon request. For determining frequencies of selected cell populations, manual gating in FlowJo was performed on cells > singlets > live cells (Zombie NIR^-^) > population of interest (as described in results). Data are presented as bar graphs using Prism (Graphpad, v10.2.3).

### Lipidomics analysis

Cells were grown in 10% exosome-depleted FBS (System Biosciences, EXO-FBSHI), 1X P/S (Corning, 30-002-Cl), DMEM (Life Technologies, 11995073) in 10cm tissue culture plates (Fisher Scientific, 08-772E), tripsinized, and pelleted. Cell pellets were stored at −80C and submitted to the University of California, Davis West Coast Metabolomics Center for downstream lipidomics analysis. Extraction of plasma lipids was based on the “Maytash’’ method (Matyash et al., 2008) which was subsequently modified. Extraction was carried out using a bi-phasic solvent system of cold methanol, methyl *tert*-butyl ether (MTBE), and water with the addition of internal standards.

The LC/QTOFMS analysis was performed using an Agilent 1290 Infinity LC system (G4220A binary pump, G4226A autosampler, and G1316C Column Thermostat) coupled to either an Agilent 6530 or an Agilent 6550 mass spectrometer equipped with an ion funnel (iFunnel). Lipids were separated on an Acquity UPLC CSH C18 column (100 x 2.1 mm; 1.7 µm) maintained at 65°C at a flow-rate of 0.6 mL/min with a 15 minute total run time. Solvent pre-heating (Agilent G1316) was used. A sample volume of 0.1uL-0.5uL for positive ion mode and 2-3uL for negative ion mode was used for the injection. Sample temperature was maintained at 4°C in the autosampler.

For the data processing, the MassHunter software (Agilent) was used, and a unique ID was given to each lipid based on its retention time and exact mass (RT_mz). This allowed the report of peak areas/heights or concentration of lipids based on the use of internal standards. Lipids were identified based on their unique MS/MS fragmentation patterns using in-house software, Lipidblast. Using complex lipid class-specific internal standards, this approach quantifies >400 lipid species including mono-, di- and triacylglycerols, glycerophospholipids, sphingolipids, cholesterol esters, ceramides, and fatty acids (Cajka and Fiehn, 2016).

Data were normalized using the mean total of identified compounds method. Univariate between-groups comparisons were conducted using independent-samples *t*-tests, with *p* values not corrected for multiple comparisons. Fold changes using medians of each group for each lipid were calculated as a measure of effect size. Data analyses were conducted in R version 4.3.2 (R Core Team, 2024).

### Statistics

Data are presented as mean plus minus standard error of the mean unless otherwise noted and were analyzed using GraphPad Prism or Microsoft Office Excel. Statistical significance was assumed at a *P* value of ≤0.05 for unpaired t tests, log-rank tests, Kruskal-Wallis test, Fisher’s exact test, chi square test, and Wilcoxon Rank Sum tests. *P* values were calculated with the unpaired t test unless indicated otherwise. For interpretation of statistical results from unpaired t tests, log-rank tests, Kruskal-Wallis test, and Wilcoxon Rank Sum tests * = *P* ≤ 0.05, ** = *P* ≤ 0.01, *** = *P* ≤ 0.001, and **** = *P* ≤ 0.0001. Outliers were detected and removed from analyses using the ROUT (Q=5%) or Grubb’s (Alpha=0.2) outlier tests in GraphPad Prism.

### Western blotting

For Supplemental Figure 7, the following immunoblotting protocol was used: Each fraction of the C-DGUC purified sEVs (37.5 μL) was mixed with 12.5 μL of 4 × Laemmli Sample Buffer (Bio-Rad) and 10% 2-Mercaptoethanol (Bio-Rad). After boiling at 95°C for 5 minutes, samples were loaded onto a 10% SDS-PAGE gel and transferred onto PVDF membrane (Bio-Rad). PVDF membranes were then blocked with 5% non-fat milk dissolved in DPBS for 1 hour. Afterwards, the membranes were probed with the following primary antibodies overnight at 4°C: anti-CD9 clone EPR2949 (1:100, Abcam, 92726), anti-CD81 clone B11 (1:100, Santa Cruz Biotechnology, sc-166029), anti-CD81 clone D4 (1:100, Santa Cruz Biotechnology cat #sc-166028), anti-CD63 (1:100, Abcam, 68418), and anti-Flotillin-1 clone D2V7J (1:500, Cell Signaling, 18634). The next day, the membranes were washed 4 times in DPBS containing 0.1% Tween (PBST), then incubated with anti-mouse IgG-HRP secondary antibody (1:1000,

Santa Cruz Biotechnology, sc-516102) or anti-rabbit IgG-HRP secondary antibody (1:1000, Thermo Fisher Scientific, A10547) for 1 hour. Membranes were once again washed 4 times with PBST, incubated for 5 minutes with Amersham ECL Prime (Cytiva, RPN2232), and imaged using an ImageQuant LAS 4000 (GE). For all other Western blots in the manuscript, the protocol we previously described in detail was used (Hendley et al., 2021).

### Cell culture

HPDE6c7 (RRID:CVCL_0P38), CFPAC-1 (ATCC, CRL-1918), Capan-1 (ATCC, HTB-79), PaTu 8988s (RRID:CVCL_1846), PaTu 8988t (RRID:CVCL_1847), PANC-1 (ATCC, CRL-1469), MIA PaCa-2 (ATCC, CRL-1420), RAW 264.7 cells, and murine PDA cell lines were grown in 10% FBS (Corning, 35011CV), 1X P/S, DMEM (Life Technologies 11995073). hTERT-HPNE (ATCC, CRL-4023) cells were grown in 75% DMEM without glucose (Sigma-Aldrich, D-5030) with additional 2 mM L-glutamine and 1.5 g/L sodium bicarbonate, 25% Medium M3 Base (Incell Corp., M300F-500), 5% FBS, 10 ng/mL human recombinant EGF (R&D Systems, 236-EG), 5.5 mM D-glucose (1g/L) (Sigma-Aldrich, G8270), 750 ng/mL puromycin (Gemini, 400-128P). BxPC-3 (ATCC, CRL-1687), Panc 03.27 (ATCC, CRL-2549), Panc 10.05 (ATCC, CRL-2547),

COLO 357 (RRID:CVCL_0221) cells were grown in 10% FBS, 1X P/S, RPMI (Corning, 15-040-CM), 0.3g/L L-Glutamine (Gibco, 21051-024). Human cell lines (except PDOs) used in this study were authenticated using ATCC’s human cell line authentication service (ATCC, 135-XV). The murine RAW 264.7 cell line was authenticated using ATCC’s murine cell line authentication service (ATCC, 137-XV). The cell line itself, a parental cell line, or a derivative of every cell line used in this study tested negative for mycoplasma (Sigma-Aldrich, MP0025). All cells used in this study were incubated at 37°C in 5% CO_2_.

Human patient-derived organoids were obtained from the Organoid D2B unit at UCSF. Human PDO#2 line was grown in DMEM/F12 [+] L-Glutamine [+] 2.438 g/L sodium bicarbonate (Gibco, 11320-033) supplemented with 1X Penicillin/Streptomycin (Corning, 30-002-Cl), 10mM HEPES (Gibco, 15630-080), 1.25mM N-Acetylcysteine (Sigma-Aldrich, A9165), 10mM Nicotinamide (Sigma-Aldrich, N0636), 100ug/mL Primocin (InvivoGen, ant-pm-05), 1X NeuroCult™ SM1 Neuronal Supplement (Stem Cells Technology, 05711), 50ng/mL hEGF (R&D Systems, 236-EG), 100ng/mL hFGF 10 (R&D Systems, 345-FG), 100ng/mL hNoggin (R&D Systems, 3344-NG), 10nM hGastrin I (Anaspec, AS-20750), 25nM RSPO-2 (Luca et al., 2020) (gift of K. Christopher Garcia), and 0.5 μM A83-01 (Cayman Chemical, 9001799). The media for human PDO line #1 is the same as #2 except without A83-01. Organoids were cultured in 40uL domes of Cultrex RGF BME, Type 2 (R&D Systems, 3533-005-02).

### Generation of primary murine pancreatic cancer cell lines

Mice were euthanized and the pancreas was dissected and placed in a 60mm dish filled with HBSS(-/-) (Gibco, 14175-095). All of the following steps were performed in a biosafety cabinet. Pancreas was placed in a 10cm tissue culture plate and washed with 10mL HBSS(-/-). Pancreas was transferred to the top lid of a 10cm tissue culture plate and minced to <1mm pieces with scissors. Pancreas was transferred to a 50mL conical tube containing 5mg Collagenase (Gibco, 17018-029) and 10mL DMEM/F12 (Gibco, 11320-033) and incubated shaking horizontally at 180rpm for 20-25 minutes at 37°C. Reaction was quenched by addition of DMEM/F12 containing 10% FBS. Cell suspension was filtered through a 100uM nylon cell strainer (Falcon, 352360) into a new 50mL conical tube. Filtered cells were spun at 1500rpm for 5 minutes. Supernatant was aspirated and cells were resuspended and plated in RPMI, 10% FBS, 1X P/S on collagen coated 10cm tissue culture plates (Corning, 354450) for P0. For P1 and after, cells were passaged to 10cm tissue culture treated plates (Fisher, 353003) and cultured in 10% FBS, 1X P/S (Corning, 30-002-Cl), DMEM (Life Technologies 11995073). Because the initial plating of cells includes multiple cell types including fibroblasts, cells were passaged to at least P4 before they were used in any experiments in this study.

### Cell culture assays

For all cell culture assays using primary murine lines, independent experiment replicates are graphed. Experiments for all cell culture assays were performed on primary cell lines below passage 30.

#### Cell titer glow assays

For cell titer glow assays, 1500 cells were seeded in 150uL 10% exosome-depleted FBS (System Biosciences, EXO-FBSHI), 1X P/S, DMEM (Life Technologies 11995073) in a 96 well plate (Corning, 3904) on Day 0. On Day 5, the cell titer glow assay was performed. The CTG reagent (Promega, G7570) was reconstituted according to the manufacturer’s instructions, diluted 1:2 in 1X PBS-/- (Gibco, 14190-144), and stored at −80°C. The diluted CTG reagent was thawed and allowed to come to RT. Media was aspirated, and 50uL diluted CTG reagent was added to each well. The plate was shaken at RT on a Corning LSE low speed orbital shaker at 45 RPM for 15 minutes, then incubated for an additional 15 minutes at RT without shaking. Luminescence was read on a Perkin Elmer Wallac 1420 Victor2 Microplate Reader.

#### Colony formation assays

For colony formation assays, 100 cells were seeded on Day 0 in a 6 well plate in 3mL media: 10% exosome-depleted FBS (System Biosciences, EXO-FBSHI), 1X P/S, DMEM (Life Technologies 11995073). On Day 11, the colonies were stained using crystal violet (Sigma, V5265) according to standard protocols. Colonies were quantified using light microscopy and defined as clusters of 50 or more adjacent cells (Crowley et al., 2016).

#### GW4869 treatment of mouse and human cell lines

6 x 10^6 CD63-GFP-expressing cells were seeded on Day 1 with either 20uM GW4869 (Santa Cruz Biotechnology, sc-218578) or vehicle (DMSO (Santa Cruz Biotechnology, sc-359032)). Twenty-four hours after seeding, conditioned media was collected, and cells were counted using a Countess II FL Automated Cell Counter (Life Technologies) for normalization. Conditioned media was filtered using a 0.22uM PES filter (Thermo Fisher Scientific, CH2225-PES), and Polyethylene glycol (PEG) M.W. 8000 (ICN Biomedicals, Inc., 194839) solution 1:5 was added to the filtered, conditioned media for precipitating vesicles overnight. Conditioned media with added PEG solution was incubated for 16.5 hours, spun at 1500g, 4°C, for 1 hour, supernatant was aspirated, and the pellet was resuspended in 200uL 1X PBS and stored at 4°C (Bittel et al., 2021). Tenfold dilutions were quantified utilizing a NanoSight NS300 and NanoSight NTA 3.4 software with a detection threshold of 2. Data are presented as the following ratio: CD63eGFP^+^ particles divided by total cell number at time of supernatant harvest. Cells were cultured in the media described in Cell Culture materials and methods section with exosome-depleted FBS (System Biosciences, EXO-FBSHI).

Addition of sEVs/exosomes to RAW 264.7 cells for polarization assessment RAW 264.7 cells were seeded at a density of 2 x 10^6^ / 100mm plate on Day 0 in media containing exosome-depleted FBS. On Day 1, media was replaced with one of five conditions: 1) media containing exosome-depleted FBS with PBS 2) 1 x 10^9^ sEVs isolated from *KPC; Smpd3^wt/wt^* PDA cell lines in media containing exosome-depleted FBS 3) 1 x 10^9^ sEVs isolated from *KPC; Smpd3^f/f^* PDA cell lines in media containing exosome-depleted FBS 4) 100ng/mL LPS (Sigma, L3024) 5) 5ng/mL recombinant murine IL-4 (PeproTech, 214-14). 4, 16, and 24 hours after addition of each condition, cells were washed with 1X PBS and lysed directly in the tissue culture plate using Buffer RLT Plus containing beta-mercaptoethanol (ß-ME) (10uL per mL). RNA was isolated and used for qPCR. Total media volume per well in a 60mm tissue culture plate was 3mL.

The sEVs used for these *in vitro* experiments were the exact same as those injected into the mouse models – see “Injection of purified sEVs/exosomes into mice” methods section for details on sEV isolation, purification, and composition.

#### IC50 determination

Gemcitabine IC50 was determined as previously described (Collisson et al., 2011). 2500 cells were seeded in a 96 well plate on day 0, treated with a range of 11 gemcitabine hydrochloride (Sigma, G6423) concentrations on day 1, and cell number was estimated using CTG assay on day 4. IC50 was calculated in Prism using [Inhibitor] vs. response -- Variable slope (four parameters) test.

### Neutral sphingomyelinase enzymatic activity assay

#### Preparation of cells grown in 10cm plates

Cells were grown as described. Media was aspirated and cells were washed 2X with 1X PBS-/- (Gibco, 14190-144) on ice. 100uL cell lysis buffer (1X cell lysis buffer (Cell Signaling, 9803S), 100 mM PMSF (Sigma-Aldrich, P7626), 1X cOmplete Protease Inhibitor Cocktail (Roche, 11697498001), and 1X PhosSTOP (Sigma-Aldrich Aldrich, 4906845001)) was added to the 10cm plate and cells were scraped using a cell lifter (Corning, 3008).

#### Preparation of organoids grown in 24 well plates

Organoids were grown as described. Organoid media was aspirated and 500uL Cell Recovery Solution (Corning, 354253) was added to each well. Cultrex dome was disrupted with a 1000uL pipette tip and collected into a 1.5mL microcentrifuge tube. The cultrex dome containing organoids was incubated for 1 hour on a rocker at 4°C to dissolve cultrex. 1000uL 1X PBS-/- was added to the microcentrifuge tube. Cells were spun at 200g for 5 min at 4C. Cells were washed 2X with 1X PBS-/-. The cell pellet was resuspended in 100uL cell lysis buffer.

#### Cell lysis procedure, protein quantification, and nSMase activity assay

Cells in lysis buffer were collected in a 1.5mL microcentrifuge tube and incubated on ice for 30 minutes. Cells were vortexed every 10 minutes during the incubation. Cells were spun at 16,873 rcf for 15 minutes at 4°C. Supernatant was collected without disturbing the pellet and stored at - 80°C.

Cell lysates were thawed on ice and quantified using the BCA Assay (Thermo Fisher Scientific, 23225). 40ug protein and two technical replicates per sample were used in the sphingomyelinase assay. The neutral sphingomyelinase assay was performed according to the manufacturer’s instruction manual (Abcam, ab138876).

### Isolation and purification of sEVs/exosomes for proteomics and small-RNA-seq

#### Conditioned Media Preparation

Cells were grown in 10% exosome-depleted FBS (System Biosciences, EXO-FBSHI), 1X P/S (Corning, 30-002-Cl), DMEM (Life Technologies 11995073) in 15cm tissue culture plates. Supernatant was collected when cells were subconfluent and spun at 400g for 10 minutes to remove dead cells. Supernatant was transferred to a new 50mL conical tube and spun at 2000g for 30 minutes to remove large cellular debris. Subsequently, supernatant was filtered using a 0.22uM filter (Millipore Sigma, SLGP033RS).

#### Isolation and Purification of sEVs/exosomes from Conditioned Media

Cushioned-Density Gradient UltraCentrifugation (C-DGUC) was employed to isolate and purify the exosomes. The filtered Conditioned Media (CM) was centrifuged on 2 mL of a 60% iodixanol cushion (Sigma-Aldrich) at 100,000 × g for 3 hours at 4°C (345776, Type 45 Ti, Beckman Coulter). Afterwards, 3 mL on the bottom of the tube was carefully extracted and loaded onto the bottom of an OptiPrep density gradient (5%, 10%, 20% w/v iodixanol), which was then centrifuged at 100,000 × g for 18 hours at 4°C (344060, SW 40 Ti, Beckman Coulter) for further purification. The next day, twelve 1 mL fractions were collected starting from the top of the tube. Fractions of interest were taken and dialyzed in DPBS (Corning) with Slide-A-Lyzer MINI Dialysis Devices (Thermo Fisher Scientific). Dialyzed samples are suitable for *in vivo* and *in vitro* studies or can be processed for further analyses.

#### Nanoparticle Tracking Analysis (NTA)

Particles in fractions of interest were subjected to size and concentration measurements by Nanosight LM14 (Malvern PANalytical) and using a 488 nm detection wavelength. Samples were prepared by diluting them 1:100 or 1:200 with DPBS. The analysis settings were optimized and standardized for each sample: the detection threshold was set at 3 and three 1-minute videos were captured and analyzed to give the mean, median, mode, and estimated concentration for each particle size. NTA 3.3 (Malvern PANalytical) software was used to collect and analyze the data.

#### Isolation of sEVs/exosomes used for proteomics analysis and small RNA-sequencing

After NTA and Western blot analyses determined that Fractions 6-9 from C-DGUC were the most concentrated samples, the fractions were pooled together. From this pool, 800 μL (200uL/1000uL per fraction) was taken and subjected to RNA extraction (see RNA isolation, cDNA library preparation, and qPCR section under MATERIALS AND METHODS for details). The extracted RNA was then subjected to small RNA sequencing. Also from this pool, 2.6 mL (650uL/1000uL per fraction) was taken, adjusted to 5mL with DPBS, and centrifuged at 100,000 × g for 5 hours at 4°C (MLS-50, Beckman Coulter, 326819). Afterwards, the supernatant was carefully removed, and the pellet was left to dry. The dried pellet was then frozen at −80°C, which was later thawed and resuspended for proteomics analysis.

### sEV/exosome proteomics

#### Sample preparation

sEV proteomics was conducted by the UC Davis Proteomics Core. sEVs were lysed with a buffer consisting of 5% SDS and 50mM triethyl ammonium bicarbonate for 1 hour on ice. The lysate’s protein concentration was measured by BCA assay (Thermo Fisher Scientific, 23225). All protein samples were subjected to clean-up / reduction / alkylation / tryptic proteolysis by using suspension-trap (ProtiFi) devices. Suspension-trap is a powerful Filter-Aided Sample Preparation (FASP) method that traps acid aggregated proteins in a quartz filter prior to enzymatic proteolysis. Here, proteins were resuspended in 50 µL SDS solubilization buffer consisting of 5% SDS, 50 mM TEAB, pH 7.55). Disulfide bonds were reduced with dithiothreitol and alkylated (in the dark) with iodoacetamide in 50mM TEAB buffer. Digestion consisted of a first addition of trypsin (1:100 enzyme: protein (wt/wt)) for 4 hours at 37 °C, followed by a boost addition of trypsin using same wt/wt ratios for overnight digestion at 37 °C. To stop the digestion, the reaction mixture was acidified with 1% trifluoroacetic acid. The eluted tryptic peptides were dried in a vacuum centrifuge and reconstituted in water with 2% acetonitrile (ACN).

#### LC-MS

Liquid chromatography was performed using an ultra-high pressure nano-flow Easy-nLC (Bruker Baltonics), at 40°C with a constant flow rate of 400 nL/min on a PepSep 150μm x 25cm C18 column (PepSep, Denmark) with 1.5μm particle size (100 Å pores) (Bruker Daltronics), and a zero dead volume spray emitter (Bruker Daltronics). Mobile phases A and B were water with 0.1% formic acid (v/v) and 80/20/0.1% ACN/water/formic acid (v/v/vol), respectively. Peptides were separated using a 60 min gradient. The gradient increased from 10 to 30% B within 30 min, followed by an increase to 40% B within 15 min and further to 57.5% B within 5 min before washing and re-equilibration. In 30 min separations, the initial 10–30% B step was 15 min, followed by a linear increase to 40% B (7.5 min) and 57.5% B (2.5 min) before washing and re-equilibration.

Mass spectrometry was performed on a hybrid trapped ion mobility spectrometry-quadrupole time of flight mass spectrometer (timsTOF Pro, (Bruker Daltonics, Bremen, Germany) with a modified nano-electrospray ion source (CaptiveSpray, Bruker Daltonics). In the experiments described here, the mass spectrometer was operated in PASEF mode. Desolvated ions entered the vacuum region through the glass capillary and deflected into the TIMS tunnel, which is electrically separated into two parts (dual TIMS). Here, the first region is operated as an ion accumulation trap that primarily stores all ions entering the mass spectrometer, while the second part performs trapped ion mobility analysis (2). Data-independent analysis (DIA) was performed on a nanoElute UHPLC coupled to a timsTOF Pro. The acquisition scheme used for DIA consisted of four 25 m/z precursor windows per 100ms TIMS scan. Sixteen TIMS scans, creating 64 total windows, layered the doubly and triply charged peptides on the m/z and ion mobility plane. Precursor windows began at 400 m/z and continued to 1200 m/z. The collision energy was ramped linearly as a function of the mobility from 63 eV at 1/K0=1.5 Vs cm−2 to 17 eV at 1/K0=0.55 Vs cm−2.

#### Data analysis

Data analysis was conducted by the UC Davis Proteomics Core facility and the UC Davis Bioinformatics Core. Files were processed with Spectronaut version 18 (Biognosys, Zurich, Switzerland) using DirectDIA analysis mode. Mass tolerance/accuracy for precursor and fragment identification was set to default settings. Global imputation was turned on. The Uniprot database of reviewed *Mus musculus* proteins (accessed 08/04/2021 from UniProt, UP000000589 and a database of 112 common laboratory contaminants (https://www.thegpm.org/crap/) were used. A maximum of two missing cleavages were allowed, the required minimum peptide sequence length was 7 amino acids, and the peptide mass was limited to a maximum of 4600 Da. Carbamidomethylation of cysteine residues was set as a fixed modification, and methionine oxidation and acetylation of protein N termini as variable modifications. A decoy false discovery rate (FDR) at less than 1% for peptide spectrum matches and protein group identifications was used for spectra filtering (Spectronaut default). Decoy database hits, proteins identified as potential contaminants, proteins identified exclusively by one site modification, and non-murine proteins were excluded from further analysis. Three samples (*KPC; Smpd3^wt/wt^* 10, *KPC; Smpd3^wt/wt^* 11, and *KPC; Smpd3^wt/wt^* 12) were excluded from the wild-type group because the total ion chromatogram deviated from the other samples due to either sample prep or instrumentation failure. Figures (volcano plot and heatmap) were created using R version 4.3.1. The heatmap was created using the R package ComplexHeatmap, version 2.16.0.

### Mice

Rag1 KO and C57BL/6J mice were purchased from Jackson Labs (Stock #002216 and Stock #000664, respectively). The following mouse strains were used: Ptf1a-Cre (Kawaguchi et al., 2002) (gift of Christopher Wright, Vanderbilt University), Kras^G12D^ (Jackson et al., 2001) (gift of David Tuveson, Cold Spring Harbor Laboratory), Trp53 flox (Marino et al., 2000), ACTB:FLPe (Jackson Labs, Stock #005703), R26R-eYFP (Jackson Labs, Stock #006148), and mTmG (Muzumdar et al., 2007). Except for flow cytometry immune profiling experiments, *KPC; Smpd3^wt/wt^*, *KPC; Smpd3^f/wt^*, and *KPC; Smpd3^f/f^* mice used in this study had a R26R-eYFP, mTmG, or no reporter allele. For end-stage time points (Figure 2A), mice were euthanized when they reached humane endpoints defined by BCS<2, hunched posture, piloerection, and decreased mobility. Mice were maintained on a mixed genetic background and genotyped by PCR or Transnetyx. The University of California at San Francisco Institutional Animal Care and Use Committee (IACUC) approved all mouse experiments in this study.

### Murine tumor model experiments

#### Subcutaneous tumor model

For subcutaneous injection, 1 x 10^^6^ cancer cells were resuspended in 100uL of 50% GFR Matrigel (Corning, 356231): 50% PBS^-/-^ and injected subcutaneously into the flank of immune-compromised Rag1 KO mice. Mice were checked for the formation of tumors, and tumor sizes were measured with a caliper. The volumes were calculated as V=ab^2^/2 (a=longest diameter; b=shortest diameter). Mice were euthanized, and tumors were dissected when they started to show signs of ulceration or exceeded 2cm in diameter per our IACUC protocol.

#### Orthotopic tumor model

For orthotopic injection of luciferase-expressing cancer cells, 3×10^^5^ cells were suspended in 50 uL of 50% Matrigel 50% PBS, and were injected orthotopically into the pancreas of immune-compromised Rag1 KO mice as previously described (Pylayeva-Gupta et al., 2012). For *in vivo* imaging, mice were administered 150 mg/kg D-luciferin (Gold Biotechnology, LUCK) by intraperitoneal injection. 10 minutes later, photons from animal whole bodies were quantified using the IVIS2000 imaging system (Xenogen). Data were analyzed using LIVINGIMAGE software (Xenogen). Animals were sacrificed, and organs were preserved at the endpoint of each study.

#### Injection of purified sEVs/exosomes into mice

1×10^8^ purified sEVs resuspended in PBS isolated from *KPC; Smpd3^wt/wt^* or *KPC; Smpd3^f/f^* end-stage polyclonal PDA cell lines using C-DGUC were injected IP into mice in a total volume of 200uL three times weekly until survival endpoint. For injected sEVs derived from *KPC; Smpd3^wt/wt^* PDA cells, isolated and dialyzed sEVs contained within fractions 6-8 were combined altogether from 6 different *KPC; Smpd3^wt/wt^* cell lines (*KPC; Smpd3^wt/wt^ 1*, *KPC; Smpd3^wt/wt^* 3, *KPC; Smpd3^wt/wt^* 5, *KPC; Smpd3^wt/wt^* 6, *KPC; Smpd3^wt/wt^* 9, and *KPC; Smpd3^wt/wt^* 11). For injected sEVs derived from *KPC; Smpd3^f/f^* PDA cells, isolated and dialyzed sEVs contained within fractions 6-8 were combined altogether from 6 different *KPC; Smpd3^f/f^* cell lines (*KPC; Smpd3^f/f^* 1, *KPC; Smpd3^f/f^* 2, *KPC; Smpd3^f/f^* 3, *KPC; Smpd3^f/f^*4, *KPC; Smpd3^f/f^* 5, and *KPC; Smpd3^f/f^* 6).

#### Injection of mice with gemcitabine

Rag1 KO mice injected with *KPC* LucFlag scrambled shRNA, *KPC* LucFlag *Smpd3* shRNA 1, and *KPC* LucFlag *Smpd3* shRNA 2 cell lines (see orthotopic tumor model MATERIALS AND METHODS section) were given 100mg/kg gemcitabine hydrochloride (Sigma, G6423) dissolved in 1X PBS or 1X PBS (for vehicle treatment) Q3Dx4 beginning on day 10 after orthotopic injection (Olive et al., 2009).

*KPC; Smpd3^wt/wt^*and *KPC; Smpd3^f/f^*mice were injected with 100mg/kg gemcitabine hydrochloride dissolved in 1X PBS or 1X PBS (for vehicle treatment) twice weekly starting at 14 weeks of age until survival endpoint.

#### Injection of mice with dextran and tomato lectin

10-11 week old mice were injected with Lycopersicon Esculentum (Tomato) Lectin (LEL, TL), DyLight 594 (Vector labs, DL-1177-1) and Fluorescein isothiocyanate–dextran (Sigma, FD2000S) as previously described (Ruscetti et al., 2020). To assess vascular permeability, 1 mg of FITC–dextran (MW 2,000,000) was administered intravenously retro-orbitally 30 minutes before sacrifice. Similarly, 100 μg of tomato lectin, DyLight 594 was injected intravenously retro-orbitally 1 hour before sacrifice to label blood vessels.

#### Isolation of murine pancreatic cells for flow cytometry

3mL cold Collagenase P (Roche, 45-11213873001) solution (4mg Collagenase P in 5mL HBSS (-/-) (Gibco, 14175-095) per mouse) was slowly injected into the common bile duct while simultaneously clamping the end of the common bile duct at its junction with the duodenum using forceps until the pancreas distended fully. The pancreas was carefully excised and transferred to a 50mL conical tube containing 2mL Collagenase P solution. Subsequently, the pancreas was digested at 37°C for 16 minutes, and then Collagenase P was quenched with an equal volume of DMEM/F12 (Gibco, 11320-033) containing 2mM EDTA. Digested pancreas was filtered using a 100uM cell strainer (Falcon, 352360) into a new 50mL conical tube. Filtered cells were spun at 1500rpm for 5 minutes. Supernatant was aspirated and the resulting cell pellet was resuspended in FACS buffer (PBS (-/-) supplemented with 2% FCS, 2mM EDTA, and 0.09% Sodium Azide).

Resuspended cells were transferred to a 96-well V bottom plate (Corning, 3894), spun at 1500rpm for 5 minutes, and supernatant was removed. Cells were subsequently blocked with 100uL/well blocking buffer (FACS buffer with 1:50 Fc Receptor Blocker (Innovex Biosciences, NB309-15)) containing 1:5000 Zombie NIR (Biolegend, 77184) on ice for 30 minutes. Blocking buffer was washed off using FACS buffer, and cells were incubated with 100uL/well fluorescent-conjugated primary antibodies recognizing extracellular epitopes diluted in FACS buffer for 30 min on ice in the dark. Primary antibodies were washed off using FACS buffer, and cells were fixed with 150uL/well fixing buffer (1 part Foxp3/Transcription Factor Fix/Perm Concentrate (4X) (TONBO Biosciences, TNB-1020-L050), 3 parts Foxp3/Transcription Factor Fix/Perm Diluent (1X)) for 10 minutes on ice in the dark. Fixing buffer was washed off with 1X Perm Buffer (Flow Cytometry Perm Buffer (10X) (TONBO Biosciences, TNB-1213-L150) diluted 1:10 in Molecular Grade Water (Cellgro, 46-000-CM)). Cells were incubated with 100uL/well fluorescent-conjugated primary antibodies recognizing intracellular epitopes diluted in FACS buffer for 30 min on ice in the dark. Primary antibodies were washed off using FACS buffer. Washed cells were resuspended in 200uL FACS buffer, transferred to FACS tubes (Falcon, 352235) through the filter cap, and analyzed on a Cytek Aurora. Primary antibodies used are listed in Supplemental Table 6.

### Generation of *Smpd3* conditional floxed allele

EUCOMM ES cells with a targeted *Smpd3* allele (Smpd3^tm1a(EUCOMM)Hmgu^) were cultured and expanded at UCSF’s Cell and Genome Engineering Core. Super-ovulated female C57BL/6 mice (4 weeks old) were mated to C57BL/6 stud males. E3.5 embryos were collected and injected with mouse ES cells into the cavity of blastocysts. Injected blastocysts were then implanted into pseudopregnant CD1 female mice. Chimeras were born and screened for germline mutations.

The *Smpd3* floxed allele was crossed to the ACTB:FLPe mouse (Jackson Labs Stock# 005703, gift of Benoit G. Bruneau) to induce recombination between FRT sites resulting in removal of the Neomycin selection cassette and the inserted lacZ reporter sequence, leaving behind the loxP-flanked *Smpd3* exon 3. Removal of the Neomycin and lacZ cassettes in *Smpd3* floxed mice was confirmed by genotyping with Transnetyx. Genotyping primers used to genotype the *Smpd3* allele were: *Smpd3* F: 5’ GGA CGT GGC CTA TCA CTG TT 3’ and *Smpd3* R: 5’ TCA GAT CTG AGA TAC TGG CC 3’. The wild-type allele PCR product is 298bp and the floxed allele is detected at 353bp.

## ACKNOWLEDGEMENTS

The authors thank Allison Y. Zhong, Trishna S. Patel, Debbie Ngow, Minju Shim, Cassandra Belair, and Karina E. Villanueva for their indispensable help with mouse colony maintenance, genotyping, and technical assistance. The authors wish to acknowledge Laura Leonhardt, Greg A. Timblin, Nilotpal Roy, Sapna Puri, Kevin M. Tharp, and Yusuf A. Hannun for helpful intellectual discussions. The authors are grateful to Rushika M. Perera and Robert H. Blelloch for reviewing and providing feedback for this manuscript. The authors thank K. Christopher Garcia and Yi Miao for providing RSPO-2 for culturing PDOs. Graphical illustrations from BioRender.com are included in this manuscript. The authors wish to acknowledge Vinh Nguyen and Vi Dang for performing bile duct injections in *KPC; Smpd3^wt/wt^* and *KPC; Smpd3^f/f^* mice used for immune profiling experiments.

This work was supported by a Richard G. Klein Fellowship in Pancreas Development, Regeneration or Cancer (to AMH). AMH was additionally supported by F32 CA221114, a Hirshberg Foundation for Pancreatic Cancer Research Seed Grant, and as a trainee on T32 CA108462. DWD was also supported by the Hirshberg Foundation for Pancreatic Cancer Research. AU was supported by a fellowship from Daiichi Sankyo Co., Ltd, Japan. AMF was supported by a Mark Foundation for Cancer Research Momentum Award. OMG was supported in part by a Shorenstein Foundation award. VMW and JPR are both supported through a Mark Foundation for Cancer Research Endeavour Program Award. VMW was additionally supported by UO1 CA250044. Organoids were generated in the Roose Organoid D2B unit, started through a UCSF PBBR TMC (Technologies, methodologies, and Cores) grant in 2018 and a gift from the Pathology department and now, in part, funded through a Mark Foundation for Cancer Research Endeavor Program grant and support from the Shorenstein Foundation (all to JPR). PDO generation was additionally supported by the Molecular Profiling in Gastrointestinal Malignancies Biorepository (IRB #: 13-12574) and the UCSF Helen Diller Family Comprehensive Cancer Biospecimen Processing Lab. This work was also supported by grants from the U.S. Department of Veterans Affairs, including VA Merit grant I01BX003928 (to RLR), and a VA Research Career Scientist Award grant IK6BX005692 (to RLR). The COMPASS study was conducted with the support of the Ontario Institute for Cancer Research (PanCuRx Translational Research Initiative) through funding provided by the Government of Ontario, the Wallace McCain Centre for Pancreatic Cancer supported by the Princess Margaret Cancer Foundation, the Terry Fox Research Institute, the Canadian Cancer Society Research Institute, and Pancreatic Cancer Canada. The study was also supported by a charitable donation from the Canadian Friends of the Hebrew University (Alex U. Soyka). Steven Gallinger is the recipient of an Investigator Award from OICR.

Work in MH’s laboratory was supported by R01 CA172045 and a pilot award from the Parker Institute for Cancer Immunotherapy (PICI). We acknowledge the Parnassus Flow Cytometry CoLab (RRID:*SCR_018206*) for assistance generating NanoSight data. Research reported here was supported in part by the DRC Center Grant NIH P30 DK063720. This work was assisted by the UCSF Center for Advanced Technologies and is supported by the Diabetes Center at UCSF. LCMS and search results were provided by the UC Davis Proteomics Core facility, and LC-MS was funded by Neil Hunter & the Howard Hughes Medical Institute. Metabolomics Workbench was supported by NIH grant U2C-DK119886 and OT2-OD030544.

## DECLARATION OF INTERESTS

AMF’s current affiliation is Liver Disease Research, Global Drug Discovery, Novo Nordisk A/S, Malov, Denmark. AU’s current affiliation is CMC Project Management Department, Technology Division, Daiichi Sankyo Co., Ltd, Tokyo, Japan. DKC is on the SAB for Immodulon Therapeutics Ltd. MH owns stocks/stock options in Viacyte, Encellin, Thymmune, EndoCrine, and Minutia. He also serves as SAB member to Thymmune and Encellin and is co-founder, SAB member, and board member for EndoCrine and Minutia.

---

**Supplemental Figure 1.**
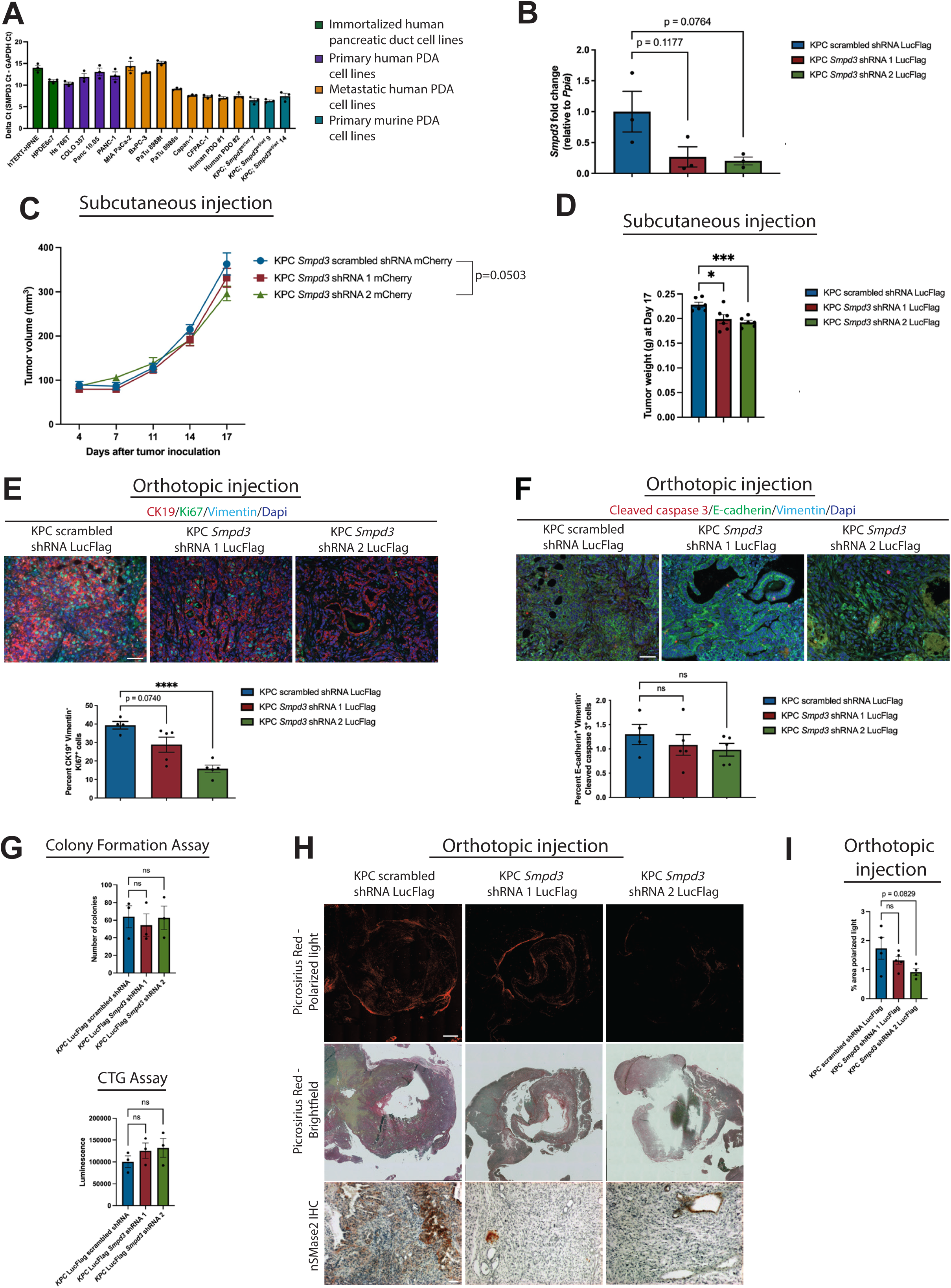
S*m*pd3 regulates proliferation of PDA cells *in vivo*. A) *SMPD3* is expressed in a panel of human pancreatic duct cells (n=2 cell lines), human PDA cells (n=12 cell lines), and mouse PDA cells (n=3 cell lines). B) qPCR demonstrates a reduction in *Smpd3* expression in KPC *Smpd3* shRNA 1 LucFlag (n=3) and KPC *Smpd3* shRNA 2 LucFlag (n=3) cell lines when compared to the KPC scrambled shRNA LucFlag (n=3) control cell line. C) Mean ± standard deviation for tumor volume of cell lines injected subcutaneously are graphed. Results of an unpaired t test comparing KPC scrambled shRNA and KPC *Smpd3* shRNA 2 cell lines at day 17 are depicted on the graph. For all cell lines, N=6 animals. D) Mean ± standard error of the mean for tumor weight at Day 17 of cell lines injected subcutaneously are graphed. For all cell lines, N=6 animals. E) Immunofluorescence images and quantification of epithelial (CK19^+^, Vimentin^-^, Dapi^+^), proliferative (Ki67^+^) pancreatic cancer cells are depicted. Average values for tumors generated using the KPC scrambled shRNA LucFlag line (n=4) are 39.33% ± 2.046%, KPC *Smpd3* shRNA 1 LucFlag line (n=5) are 28.88% ± 4.096%, and KPC *Smpd3* shRNA 2 LucFlag line (n=5) are 15.83% ± 1.960. Scale bar is 50uM. F) Immunofluorescence images and quantification of epithelial (E-cadherin^+^, Vimentin^-^, Dapi^+^), apoptotic (Cleaved caspase 3^+^) pancreatic cancer cells are depicted. Average values for tumors generated using the KPC scrambled shRNA LucFlag cell line (n=4) are 1.299% ± 0.2075%, KPC *Smpd3* shRNA 1 LucFlag cell line (n=5) are 1.082% ± 0.2126%, and KPC *Smpd3* shRNA 2 LucFlag cell line (n=5) are 0.9832% ± 0.1307. Scale bar is 50uM. G) CTG assay or colony formation assay results for the indicated cell lines are depicted. H) Reduction of *Smpd3* in epithelial pancreatic cancer cells modestly reduces fibrosis. IHC demonstrates loss of nSMase2 in epithelial pancreatic cancer cells in tumors generated using the KPC *Smpd3* shRNA 1 LucFlag and KPC *Smpd3* shRNA 2 LucFlag cell lines. One scale bar (1000uM) on the picrosirius red image is representative for all picrosirius red and corresponding brightfield images. One scale bar (200uM) on the nSMase2 IHC image is representative for all nSMase2 IHC images. I) Quantification of polarized light using the picrosirius red stained images is depicted. Average values for tumors generated using the KPC scrambled shRNA LucFlag cell line (n=4) are 1.734% ± 0.3760%, KPC *Smpd3* shRNA 1 LucFlag cell line (n=5) are 1.319% ± 0.1328%, and KPC *Smpd3* shRNA 2 LucFlag cell line (n=4) are 0.9188% ± 0.1124.

**Supplemental Figure 2.**
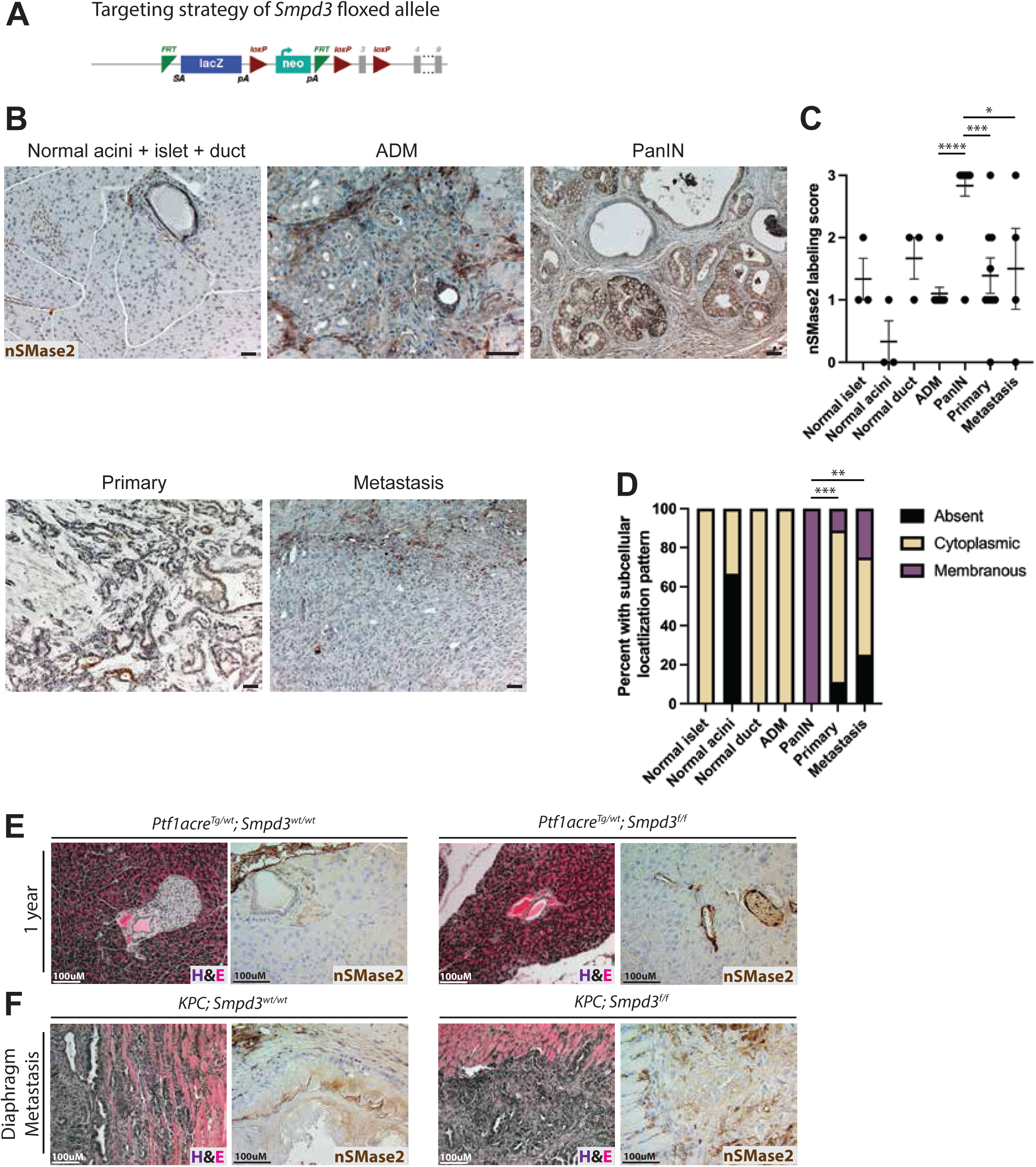
nSMase2 expression is upregulated in PanIN. A) Schematic details the construct design for the *Smpd3* floxed mouse. B-C) For examination of nSMase2 expression in the indicated normal murine pancreatic cell types, nSMase2 IHC sections from 3 C57BL/6J mice were scored. Preneoplasia and neoplasia lesions were scored from KPC mice (n=10 animals scored for acinar-to-ductal metaplasia (ADM), n=12 animals scored for PanIN, n=10 animals scored for primary tumor, and n=4 animals scored for metastasis). Scale bars for IHC images are 40 uM. nSMase2 expression was scored on a scale of 0-3 and is depicted in (C). The scoring system used is 0=absent, 1=low, 2=medium, 3=high. Values depicted are representative for individual animals. D) Subcellular localization of nSMase2 was scored using the same slides and animals used for expression scoring in Supplemental Figure 2B-C. Results from Chi-squared tests using a 2 x 3 table are shown. E-F) H&E and IHC for nSMase2 images are displayed for the indicated genotypes.

**Supplemental Figure 3.**
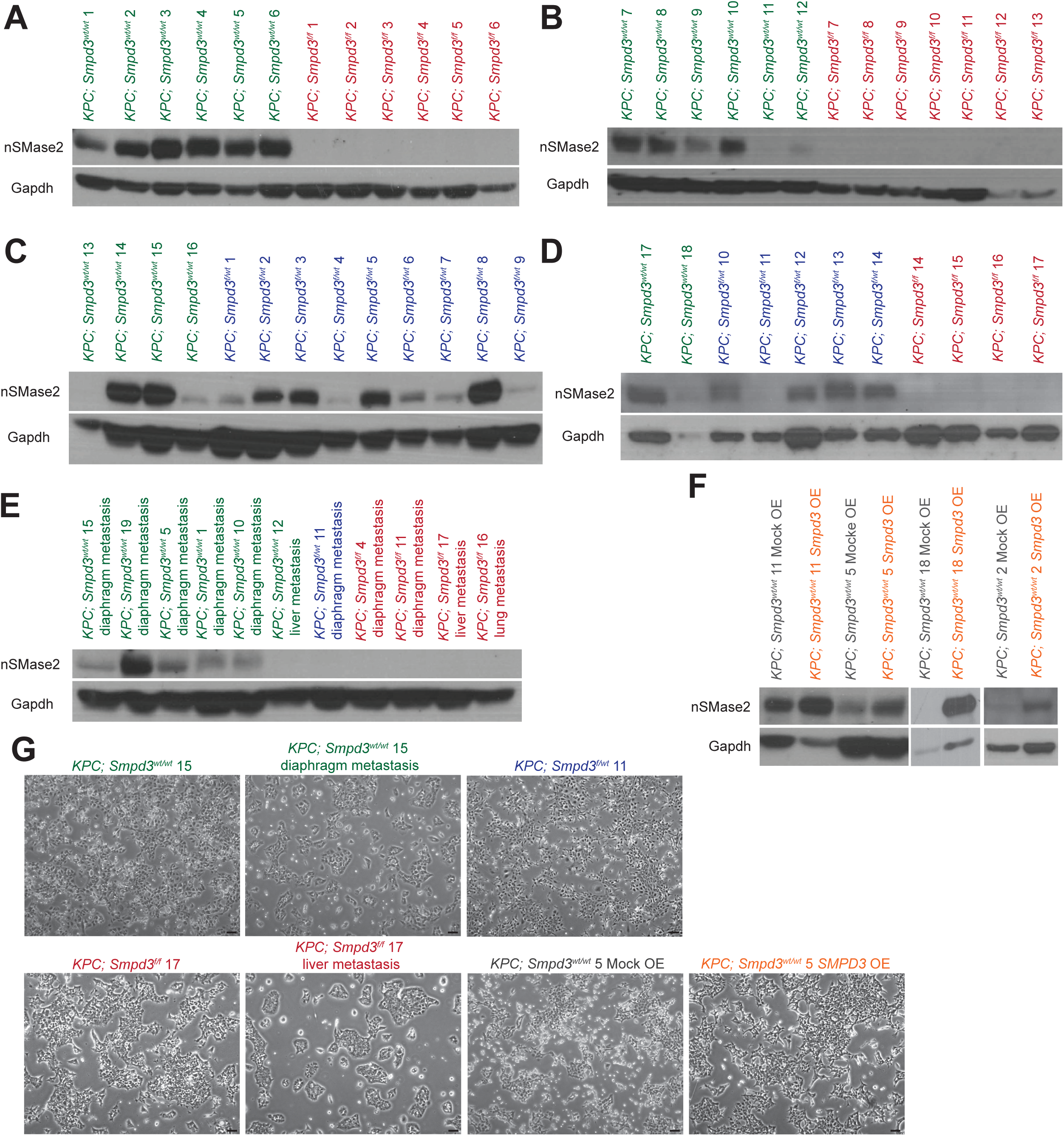
Characterization of the *KPC; Smpd3^wt/wt^*, *KPC; Smpd3^f/wt^*, *KPC; Smpd3^f/f^*, *KPC; Smpd3^wt/wt^* Mock OE, and *KPC; Smpd3^wt/wt^ Smpd3* OE cell lines. A-F) Western blot depicts nSMase2 expression levels in *KPC; Smpd3^wt/wt^*, *KPC; Smpd3^f/wt^*, *KPC; Smpd3^f/f^*, *KPC; Smpd3^wt/wt^* Mock OE, and *KPC; Smpd3^wt/wt^ Smpd3* OE cell lines. G) Pictures of cell lines taken immediately before RNA isolation for RNA sequencing are shown. Scale bars are 100uM.

**Supplemental Figure 4.**
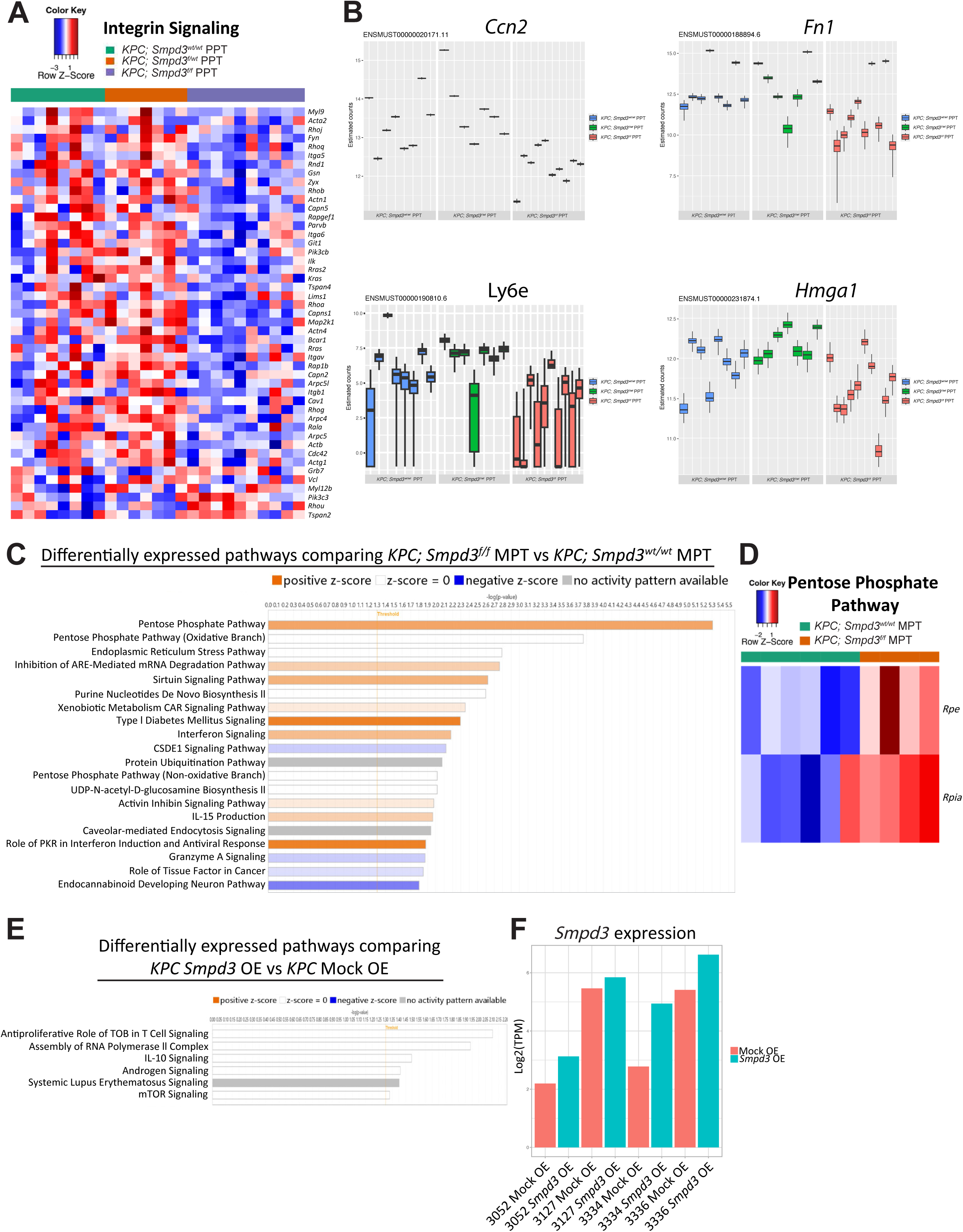
Analysis of RNA sequencing data from murine polyclonal pancreatic cancer cell lines. A) Heat map showing Integrin Signaling in all three PPT cell lines. B) Estimated counts for select genes identified from Sleuth comparison of polyclonal cell lines from end-stage PPTs of *KPC; Smpd3^wt/wt^* (n=8), *KPC; Smpd3^f/wt^* (n=7), and *KPC; Smpd3^f/f^* (n=10) mice are graphed. Each box plot represents the distribution of estimated counts quantified by kallisto with 100 bootstrap samples. For each gene, the most significant protein-coding isoforms are shown. C) The top twenty pathways from IPA comparing *KPC; Smpd3^f/f^* MPT vs *KPC; Smpd3^wt/wt^*MPT are shown. D) A heat map including significantly differentially regulated molecules in the Pentose Phosphate Pathway are shown for *KPC; Smpd3^wt/wt^* MPT and *KPC; Smpd3^f/f^* MPT groups. E) The six differentially regulated pathways when comparing *KPC Smpd3* OE vs *KPC* Mock OE are shown. F) Log of transcript per million (TPM) values for *Smpd3* gene are graphed for *KPC* Mock OE and *KPC Smpd3* OE sequenced cell lines.

**Supplemental Figure 5.**
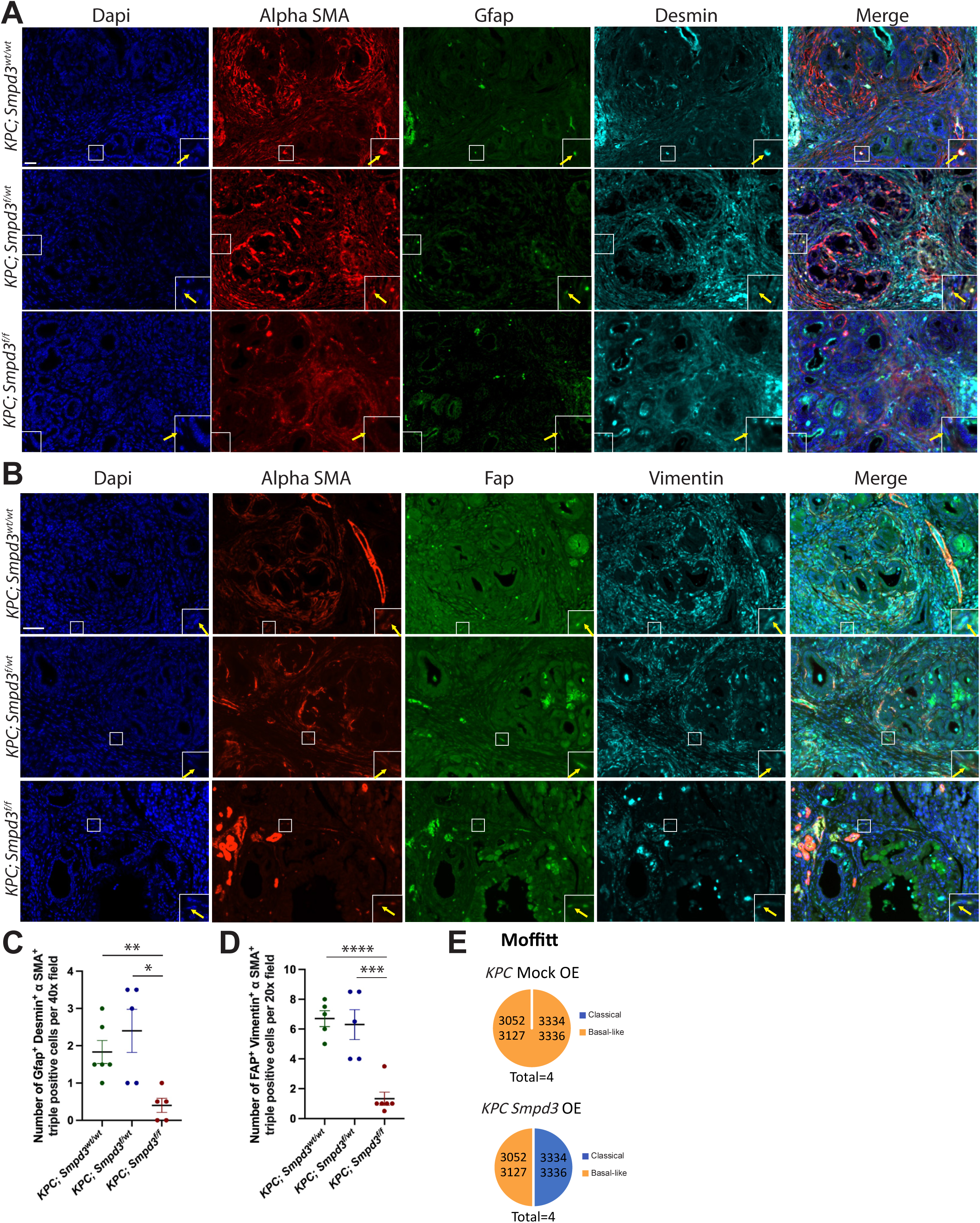
K*P*C*; Smpd3^f/f^* pancreata display significantly fewer activated stellate cells and fibroblasts when compared to both *KPC; Smpd3^f/wt^*and *KPC; Smpd3^wt/wt^* pancreata. A) Immunofluorescence images depict co-labeling with alpha SMA, Gfap, Desmin, and Dapi in the pancreas of 10–11-week-old mice. For *KPC; Smpd3^wt/wt^*, n=6 animals, *KPC; Smpd3^f/wt^*, n=5 animals, and *KPC; Smpd3^f/f^*, n=5 animals. Yellow arrows in insets point to examples of quadruple positive cells that were quantified in Panel C. Scale bar is 20uM. B) Immunofluorescence images depict co-labeling with alpha SMA, Fap, Vimentin, and Dapi in the pancreas of 10–11-week-old mice. Yellow arrows in insets point to examples of quadruple positive cells that were quantified in Panel D. For *KPC; Smpd3^wt/wt^*, n=5 animals, *KPC; Smpd3^f/wt^*, n=5 animals, *KPC; Smpd3^f/f^*, n=6 animals. Scale bar is 50uM. C) Quantification of alpha SMA, Gfap, Desmin, and Dapi quadruple positive cells is depicted. D) Quantification of alpha SMA, Fap, Vimentin, and Dapi quadruple positive cells is depicted. E) Pie charts depict Moffitt classification PDA subtypes for *KPC* Mock OE and *KPC Smpd3* OE cell lines.

**Supplemental Figure 6.**
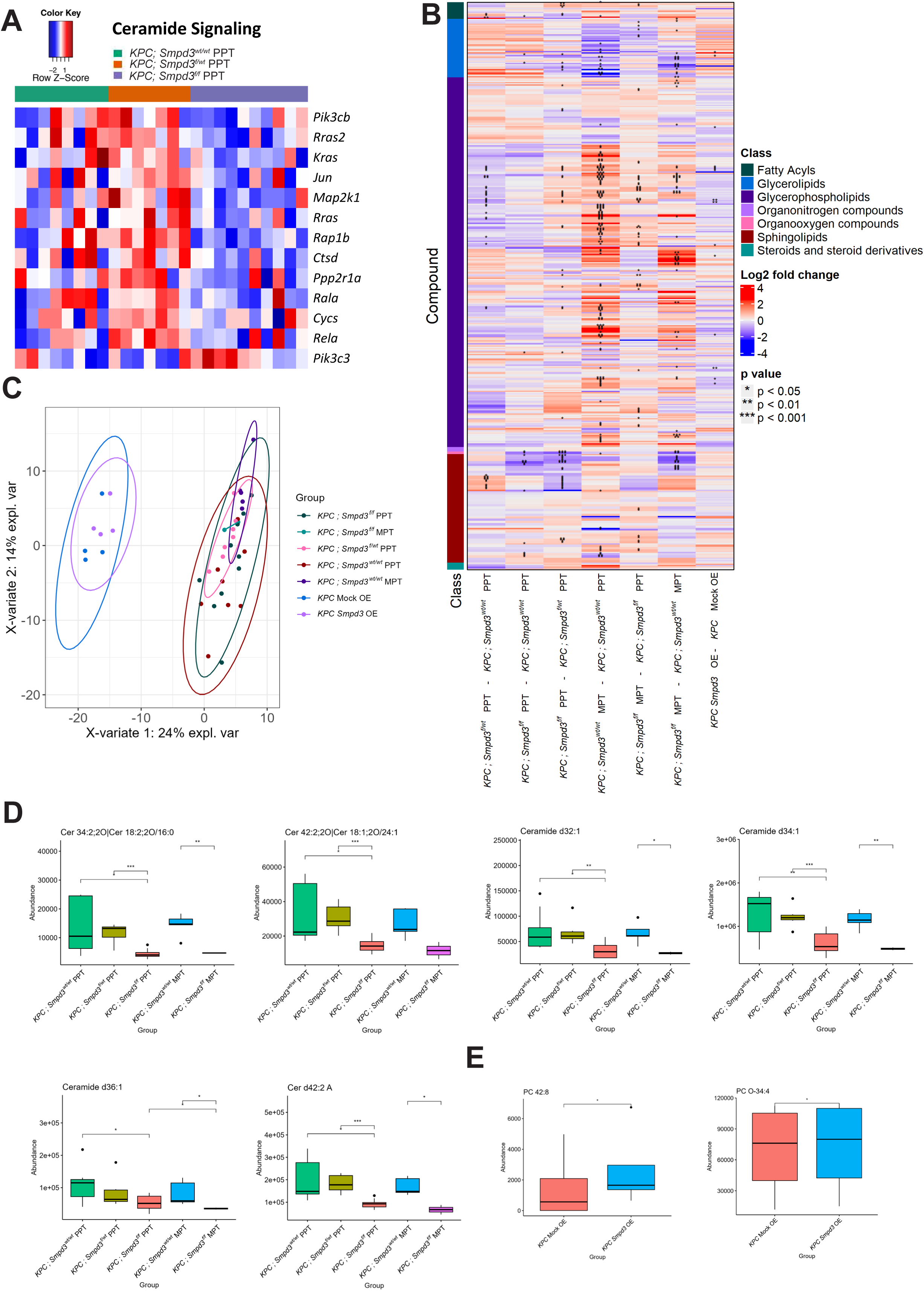
Analysis of lipidomics data from murine polyclonal pancreatic cancer cell lines. A) Heat map shows IPA Ceramide Signaling Pathway in all three PPT cell lines. B) Heat map shows differentially expressed lipids in comparisons amongst our murine polyclonal pancreatic cancer cell lines. Groups compared are listed as A-B where depicted upregulation or downregulation of a lipid would be in A when compared to B. C) Principal component analysis of our polyclonal murine pancreatic cancer cell lines based on lipid composition is depicted. D) Box plots showing expression of selected ceramide species in our *KPC; Smpd3^wt/wt^*PPT, *KPC; Smpd3^f/wt^* PPT, *KPC; Smpd3^f/f^* PPT, *KPC; Smpd3^wt/wt^* MPT, and *KPC; Smpd3^f/f^* MPT cell lines. E) Box plots showing expression of selected phosphatidylcholine species in our *KPC* Mock OE and *KPC Smpd3* OE cell lines.

**Supplemental Figure 7.**
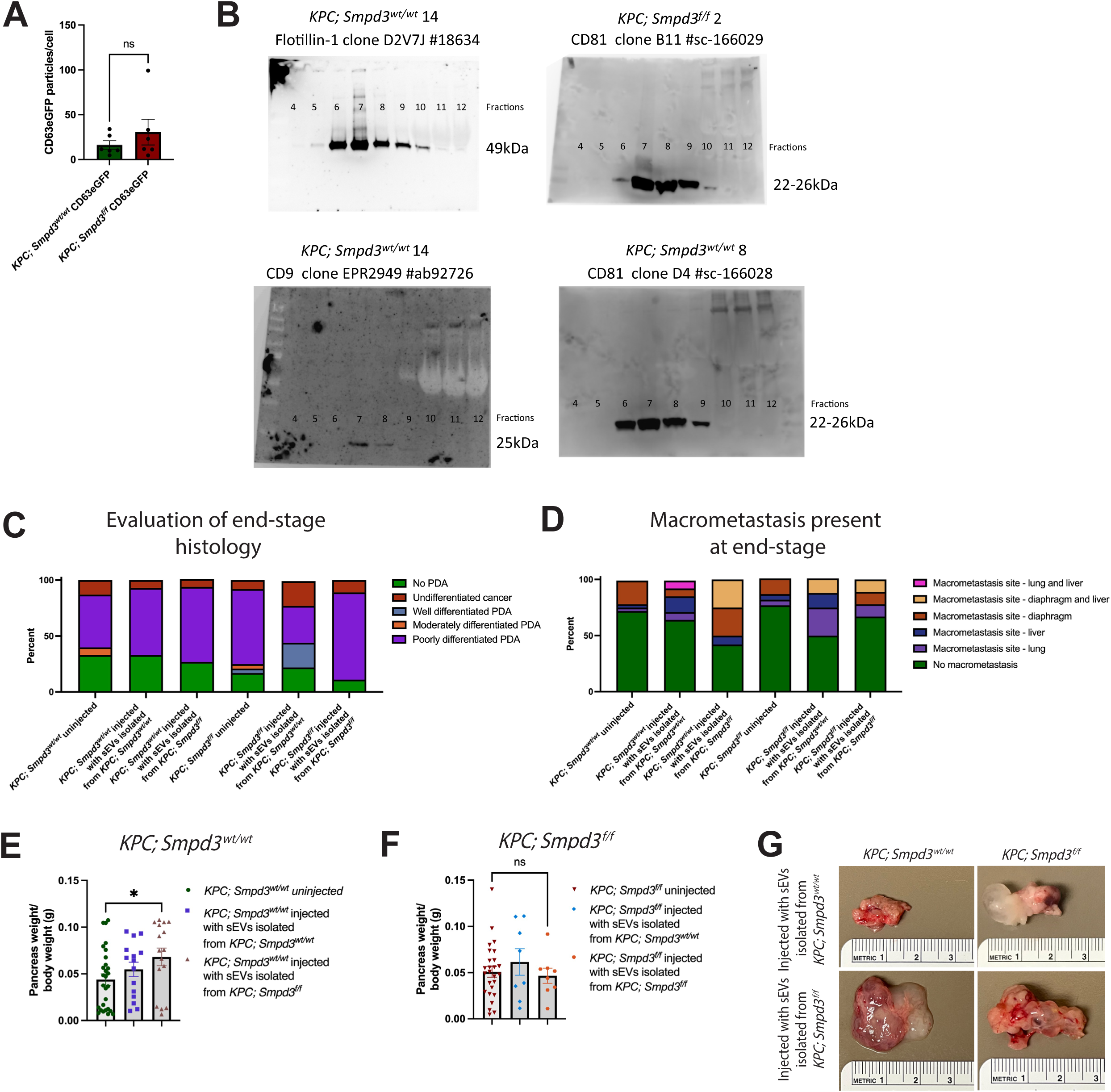
Characterization of exosomes isolated from *KPC; Smpd3^wt/wt^* PPT and *KPC; Smpd3^f/f^* PPT cell lines. A) CD63eGFP particles per cell are graphed for *KPC; Smpd3^wt/wt^* CD63eGFP cell lines (n=6) and *KPC; Smpd3^f/f^* CD63eGFP cell lines (n=6). B) Fractions were subjected to WB for sEV markers to determine the fractions that exosomes are contained within using the C-DGUC exosome isolation protocol. Expected band sizes are listed for each marker. C) The percentage of animals having PDA at end-stage in addition to the differentiation status of primary pancreatic tumors at end-stage is graphed for *KPC; Smpd3^wt/wt^* uninjected mice (n=30), *KPC; Smpd3^wt/wt^* mice injected with sEVs isolated from *KPC; Smpd3^wt/wt^* PPT cell lines (n=15), *KPC; Smpd3^wt/wt^* mice injected with sEVs isolated from *KPC; Smpd3^f/f^* PPT cell lines (n=15), *KPC; Smpd3^f/f^* uninjected mice (n=24), *KPC; Smpd3^f/f^* mice injected with sEVs isolated from *KPC; Smpd3^wt/wt^* PPT cell lines (n=9), and *KPC; Smpd3^f/f^* mice injected with sEVs isolated from *KPC; Smpd3^f/f^* PPT cell lines (n=9). Values plotted for uninjected mice are the same as those in Figure 2B. D) The percentage of animals having macrometastasis at end-stage in addition to the site of macrometastasis at end-stage is depicted for *KPC; Smpd3^wt/wt^* uninjected mice (n=29), *KPC; Smpd3^wt/wt^*mice injected with sEVs isolated from *KPC; Smpd3^wt/wt^* PPT cell lines (n=14), *KPC; Smpd3^wt/wt^* mice injected with sEVs isolated from *KPC; Smpd3^f/f^* PPT cell lines (n=12), *KPC; Smpd3^f/f^* uninjected mice (n=22), *KPC; Smpd3^f/f^* mice injected with sEVs isolated from *KPC; Smpd3^wt/wt^* PPT cell lines (n=8), and *KPC; Smpd3^f/f^* mice injected with sEVs isolated from *KPC; Smpd3^f/f^* PPT cell lines (n=9). Values plotted for uninjected mice are the same as those in Figure 2C. E) Pancreas weight/body weight is graphed for the indicated groups. Average values for uninjected *KPC; Smpd3^wt/wt^* are 0.04396 ± 0.006112 (n=28), *KPC; Smpd3^wt/wt^* injected with exosomes isolated from *KPC; Smpd3^wt/wt^* are 0.05477 ± 0.007678 (n=15), and *KPC; Smpd3^wt/wt^*injected with exosomes isolated from *KPC; Smpd3^f/f^* are 0.06809 ± 0.009409 (n=16). F) Pancreas weight/body weight is graphed for the indicated groups. Average values for uninjected *KPC; Smpd3^f/f^* are 0.05085 ± 0.005912, *KPC; Smpd3^f/f^* injected with exosomes isolated from *KPC; Smpd3^wt/wt^* are 0.06149 ± 0.01434, and *KPC; Smpd3^f/f^* injected with exosomes isolated from *KPC; Smpd3^f/f^*are 0.04654 ± 0.008279. G) Pancreas images at dissection for the indicated groups are depicted.

**Supplemental Figure 8.**
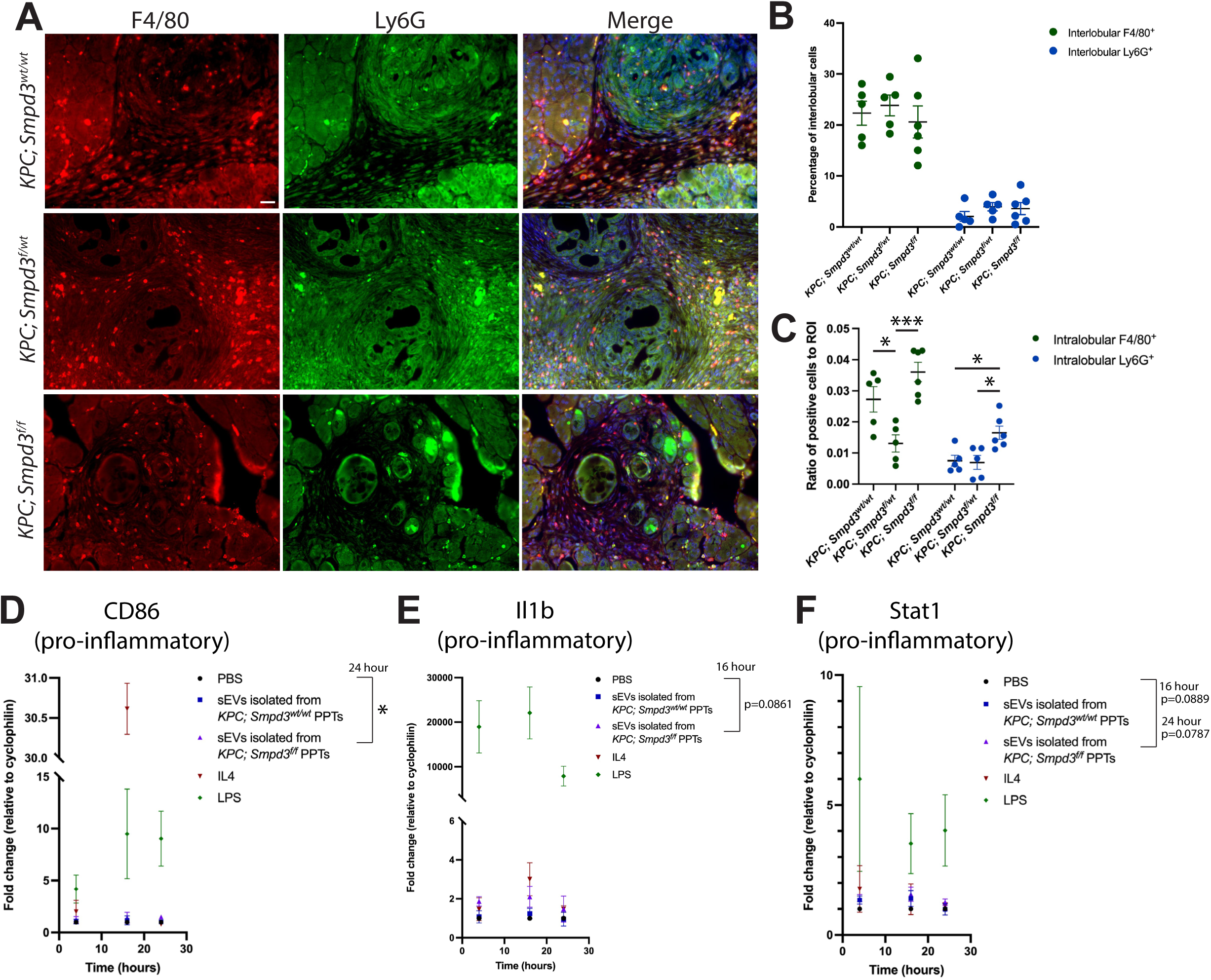
Functional effects of PDA cell sEVs on macrophages. A) Immunofluorescence images depict co-labeling with F4/80, Ly6G, and Dapi at 10-11 weeks of age. For *KPC; Smpd3^wt/wt^*, n=5 animals, *KPC; Smpd3^f/wt^*, n=5 animals, and *KPC; Smpd3^f/f^*, n=6 animals. Scale bar is 20uM. B) Quantification of interlobular F4/80 and Ly6G in *KPC; Smpd3^wt/wt^*, *KPC; Smpd3^f/wt^*, and *KPC; Smpd3^f/f^* pancreata at 10-11 weeks of age is depicted. C) Quantification of intralobular F4/80 and Ly6G in *KPC; Smpd3^wt/wt^*, *KPC; Smpd3^f/wt^*, and *KPC; Smpd3^f/f^* pancreata at 10-11 weeks of age is depicted. D-F) RNA expression fold change in RAW 264.7 cells, with PBS treated normalized to 1, for each gene at the indicated time point and treatment condition is depicted. Results of unpaired t tests between groups shown at the time point written are depicted.

**Supplemental Figure 9.**
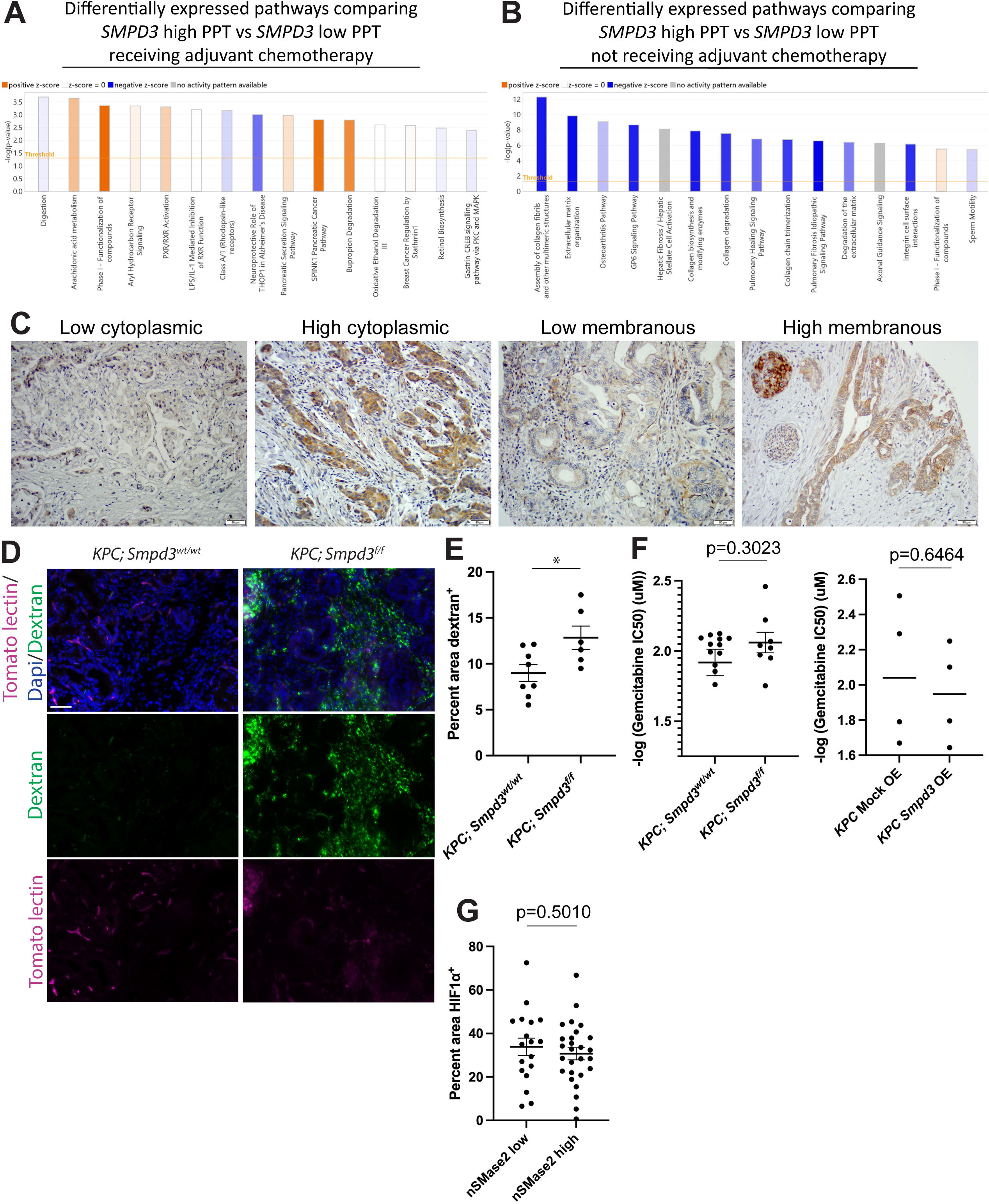
Pathways analysis of pancreatic nSMase2 expression in PDA patients based on treatment with chemotherapy in addition to the effect of nSMase2 expression on chemosensitivity and vasculature integrity. A) IPA bar chart results show the top 15 deregulated pathways when comparing PDA patients with high *SMPD3* expression in treatment naïve PPTs vs low *SMPD3* expression in treatment naïve PPTs receiving adjuvant chemotherapy. The threshold line represents statistical significance. B) IPA bar chart results show the top 15 deregulated pathways when comparing PDA patients with high *SMPD3* expression in treatment naïve PPTs vs low *SMPD3* expression in treatment naïve PPTs not receiving adjuvant chemotherapy. C) Representative images used to score the TMA for nSMase2 low and nSMase2 high membrane and cytoplasmic labeling. D) Images depict dextran and tomato lectin, a marker of blood vessels in mice, immunofluorescence at 10-11 weeks of age in *KPC; Smpd3^wt/wt^* and *KPC; Smpd3^f/f^* pancreases. E) Percent pancreatic neoplastic area per field occupied by dextran is graphed for *KPC; Smpd3^wt/wt^* (n=8) and *KPC; Smpd3^f/f^* (n=6) mice. F) Negative log of gemcitabine IC50 for *KPC; Smpd3^wt/wt^*, *KPC; Smpd3^f/f^*, *KPC* Mock OE, and *KPC Smpd3* OE cell lines is graphed. G) Quantification of total HIF1α in nSMase2 high and nSMase2 low primary pancreatic tumors of PDA patients is depicted.

Supplemental Table 1. List of polyclonal murine pancreatic cancer cell lines.

Supplemental Table 2. Analyses conducted using RNA sequencing data of murine pancreatic cancer cell lines. Results from differentially expressed genes, IPA, Sleuth analysis comparing all three PPT groups, and transcriptional subtyping analysis are shown.

Supplemental Table 3. Lipidomics analysis of polyclonal murine pancreatic cancer cell lines.

Supplemental Table 4. Exosomal small RNA-seq and proteomics analysis. Cell lines used for exosomal small RNA-Seq and proteomics are listed. Differentially expressed exosomal microRNAs and proteins are listed. IPA of proteomics data is shown.

Supplemental Table 5. Analyses conducted using RNA sequencing data of PDA patient treatment naïve PPTs. Results from differentially expressed genes and IPA are shown.

Supplemental Table 6. Table lists primary antibodies and qPCR sequences used in this study.

